# Dual histone methylation reader ZCWPW2 links histone methylation to initiation of meiotic recombination

**DOI:** 10.64898/2026.05.22.727067

**Authors:** Shenli Yuan, Shiyu Wang, Ziyou Bao, Ziqi Wang, Kang Shangguan, Yelian Yan, Yajun Shi, Xu Feng, Chengqi Huang, Mengrui Fang, Han Zhao, Shigang Zhao, Hongbin Liu, Zi-Jiang Chen, Tao Huang

## Abstract

Meiotic homologous recombination initiates with the formation of programmed DNA double-strand breaks (DSBs) by complexes comprising SPO11 and accessory proteins at discrete sites called recombination hotspots. In mammals, PRDM9-dependent H3K4me3 and H3K36me3 define recombination hotspots, but how these epigenetic characteristics determine the physiological DSB formation remains unknown. Here we show that dual histone methylation reader ZCWPW2 can recognize H3K4me3 and H3K36me3 marks in testis. The binding activity of ZCWPW2 to histone methylation is dependent on PRDM9 function. Moreover, we find the epigenetic writer-reader axis PRDM9-ZCWPW2 is essential for DSB formation at meiotic recombination hotspots, which may be partly explained by the finding that ZCWPW2 can physically interact with HORMAD1, IHO1, MEI4, and REC114. Finally, the absence of ZCWPW2 leads to disrupted chromosomal synapsis and recombination, thereby obstructing meiotic progression. Taken together, our findings provide new insights into how histone modifications and their associated regulatory proteins collectively regulate meiotic homologous recombination initiation.

## Introduction

During meiosis, the genome undergoes a single round of DNA replication followed by two successive cell divisions. During prophase I of meiosis, homologous chromosomes undergo pairing, synapsis, recombination, and segregation[1]. Meiotic homologous recombination not only enables genetic exchange between homologous chromosomes, but also ensures their accurate segregation; defects in this process can lead to severe gametogenic failure [2, 3]. Meiotic recombination is initiated by programmed DNA double-strand breaks (DSBs), which are introduced at specific genomic loci by an evolutionarily conserved topoisomerase-like complex composed of SPO11 and its accessory factors [4–9]. The resulting DNA ends are subsequently processed by the MRN (MRE11–RAD50–NBS1) complex, which resects the 5′ ends to generate 3′ single-stranded DNA (ssDNA) overhangs [10, 11]. These ssDNA intermediates are initially coated by the heterotrimeric RPA complex and are then handed over to the RecA-family recombinases RAD51 and DMC1 through BRCA2-mediated loading [12, 13]. RAD51- and DMC1-coated nucleoprotein filaments search for homologous templates and promote strand invasion, thereby generating recombination intermediates that are ultimately resolved as crossover or non-crossover products through the coordinated action of multiple repair factors[1, 14].

The number and distribution of homologous recombination events on chromosomes are tightly regulated both temporally and spatially. Meiotic homologous recombination is initiated by the formation of programmed DSBs, whose locations are not randomly distributed but rather preferentially occur at recombination hotspots[15]. In yeast, recombination hotspots are predominantly localized to nucleosome-depleted gene promoter regions, which are typically enriched in H3K4me3 modifications[16]. In sharp contrast to yeast, recombination hotspots in mammals are generally not situated in gene promoter regions. In 2010, three independent studies concurrently confirmed that *Prdm9* is the key gene governing the localization of recombination hotspots in mammals[17–19]. Subsequent studies have further demonstrated that PRDM9 is involved in determining the location of recombination hotspots in humans, non-human primates, rodents, and ruminants[20, 21].

In yeast, H3K4me3 is deposited primarily by the COMPASS complex, which contains the catalytic subunit Set1 and multiple accessory proteins[22]. On the basis of this chromatin landscape, Blat et al. proposed the “loop-axis” model in 2002 to explain how histone modifications and higher-order chromosome organization cooperate to specify meiotic DSB sites[23]. This model was subsequently validated by two independent studies in 2012. Mechanistically, these studies showed that Spp1, a COMPASS subunit, acts as a molecular bridge: through its PHD finger, Spp1 recognizes H3K4me3 near recombination hotspots, while simultaneously interacting with Mer2, an essential component of the meiotic DSB machinery[24, 25]. In this way, Spp1 couples an epigenetic chromatin mark to the execution of programmed DSB formation. Whether an analogous reader-mediated mechanism operates in mammals has remained unclear.

PRDM9 is a meiosis-specific histone methyltransferase whose central PR/SET domain catalyzes the deposition of H3K4me3 and H3K36me3. Its highly polymorphic C-terminal zinc-finger array determines its DNA-binding specificity and thereby defines the genomic locations of recombination hotspots[26]. In *Prdm9*-deficient mice, DSBs are redistributed from canonical hotspots to gene promoter regions, resulting in defective DSB repair, impaired meiotic recombination, and sterility[27]. Although PRDM9-dependent H3K4me3 and H3K36me3 have been firmly implicated in hotspot specification in both mice and humans, the downstream factors that read these histone modifications and couple them to DSB formation remain poorly defined[28]. In mice, CXXC1 (also known as CFP1), a homolog of yeast Spp1, contains an H3K4me3-binding PHD domain and has been reported to interact with both PRDM9 and IHO1[29, 30]. However, genetic ablation of *Cxxc1* does not disrupt meiotic DSB formation[31], suggesting that additional mammalian meiosis-specific factors may function to bridge hotspot-associated histone methylation with the DSB machinery.

Recent evolutionary analyses have shown that PRDM9 co-evolved with ZCWPW1 and ZCWPW2 in mammals, and that all three genes are highly expressed in leptotene and zygotene spermatocytes[32]. In corn snakes, PRDM9 directs some recombination events through its binding sites, while others occur near promoter-like regions. ZCWPW2, an epigenetic reader of histone methylation marks, may facilitate this dual usage by altering its binding affinity for H3K36me3. This finding reveals that in corn snakes, recombination hotspots involve a tug-of-war between PRDM9 binding sites and promoter-like features, which is not mutually exclusive[33]. ZCWPW2 contains both a CW domain and a PWWP domain, two modules predicted to recognize H3K4me3 and H3K36me3, respectively, raising the possibility that it may function as a dual histone methylation reader during meiosis[34]. We therefore sought to determine whether ZCWPW2 acts downstream of PRDM9 to interpret hotspot-associated histone methylation and regulate meiotic DSB formation, homologous chromosome synapsis, and recombination progression.

Here, we show that ZCWPW2 is highly and dynamically expressed in mouse testes, exhibiting stage-specific nuclear localization during meiotic prophase I and acting as a dual histone reader recognizing both H3K4me3 and H3K36me3. Genomically, ZCWPW2 preferentially occupies PRDM9 - dependent meiotic hotspots, and its chromatin binding is largely dependent on PRDM9-mediated H3K4me3 and H3K36me3 deposition. *Zcwpw2* knockout leads to complete male infertility, severe meiotic arrest at a pachytene-like stage, defective autosomal synapsis, persistent DSB signals, and loss of crossover formation. Functional inactivation of either the CW or PWWP domain fully recapitulates the meiotic failure phenotype of *Zcwpw2* knockout mice. Moreover, ZCWPW2 is indispensable for faithful DSB positioning at PRDM9-defined hotspots; its absence abolishes canonical hotspot DSBs and triggers ectopic DSB formation at promoter regions. Furthermore, ZCWPW2 physically interacts with core axis and DSB machinery components including HORMAD1, IHO1, MEI4, and REC114. Together, these findings establish ZCWPW2 as a critical chromatin effector downstream of PRDM9 that couples histone methylation recognition to accurate meiotic DSB patterning, homologous synapsis, and recombination progression.

## Results

### The histone methylation reader ZCWPW2 binds to sites with both H3K4me3 and H3K36me3 marks

We initially investigated ZCWPW2 expression across mouse tissues. As depicted in Fig. S1A, the ZCWPW2 protein demonstrates remarkably high expression in testes. Immunoblotting analysis of mouse testes at multiple developmental stages showed that ZCWPW2 protein levels gradually increased from postnatal day 6 (PD6) and exhibited high expression at PD14, coinciding with the entry of the first wave of spermatocytes into the leptotene-to-zygotene stages of meiosis (Fig.S1B). Also, the expression pattern of ZCWPW2 at multiple developmental testes is similar with PRDM9 and ZCWPW1 (Fig.S1B). These results imply that ZCWPW2 may participate in the early stages of spermatogenesis.

To clarify the subcellular localization of ZCWPW2 during spermatogenesis, we constructed a FLAG-tagged *Zcwpw2* knock-in mice (Fig.S1C). Subsequently, we performed co-immunofluorescence staining of FLAG with γ-H2AX, a well-characterized marker of DNA damage response, to distinguish germ cells at distinct developmental stages. Subcellular distribution analysis revealed that ZCWPW2 displays highly dynamic localization patterns during spermatogenesis, with prominent enrichment within the nuclei of spermatocytes and elongating spermatids. During leptotene, ZCWPW2 shows a diffuse pan-nuclear localization, which progressively assembles into distinct foci from late zygotene stage onward. Notably, in leptotene and zygotene spermatocytes, ZCWPW2 is excluded from heterochromatic chromocenter structures. During the pachytene and diplotene stages of meiosis, ZCWPW2 exclusively accumulates at the XY body. Upon completion of meiosis, nuclear accumulation of ZCWPW2 is reinstated in elongating spermatids (Fig.1A and S1C). Immunofluorescence staining of wild-type testicular sections with an anti-ZCWPW2 antibody revealed an identical localization pattern of ZCWPW2, which was consistent with that observed in the testes from FLAG-tagged *Zcwpw2* knock-in mice (Fig.1A and S1D). Collectively, these findings indicate that ZCWPW2 is highly enriched in spermatocytes and exhibits dynamic nuclear localization during meiosis, implying its role as a chromatin regulator in spermatogenesis.

**Figure 1.**
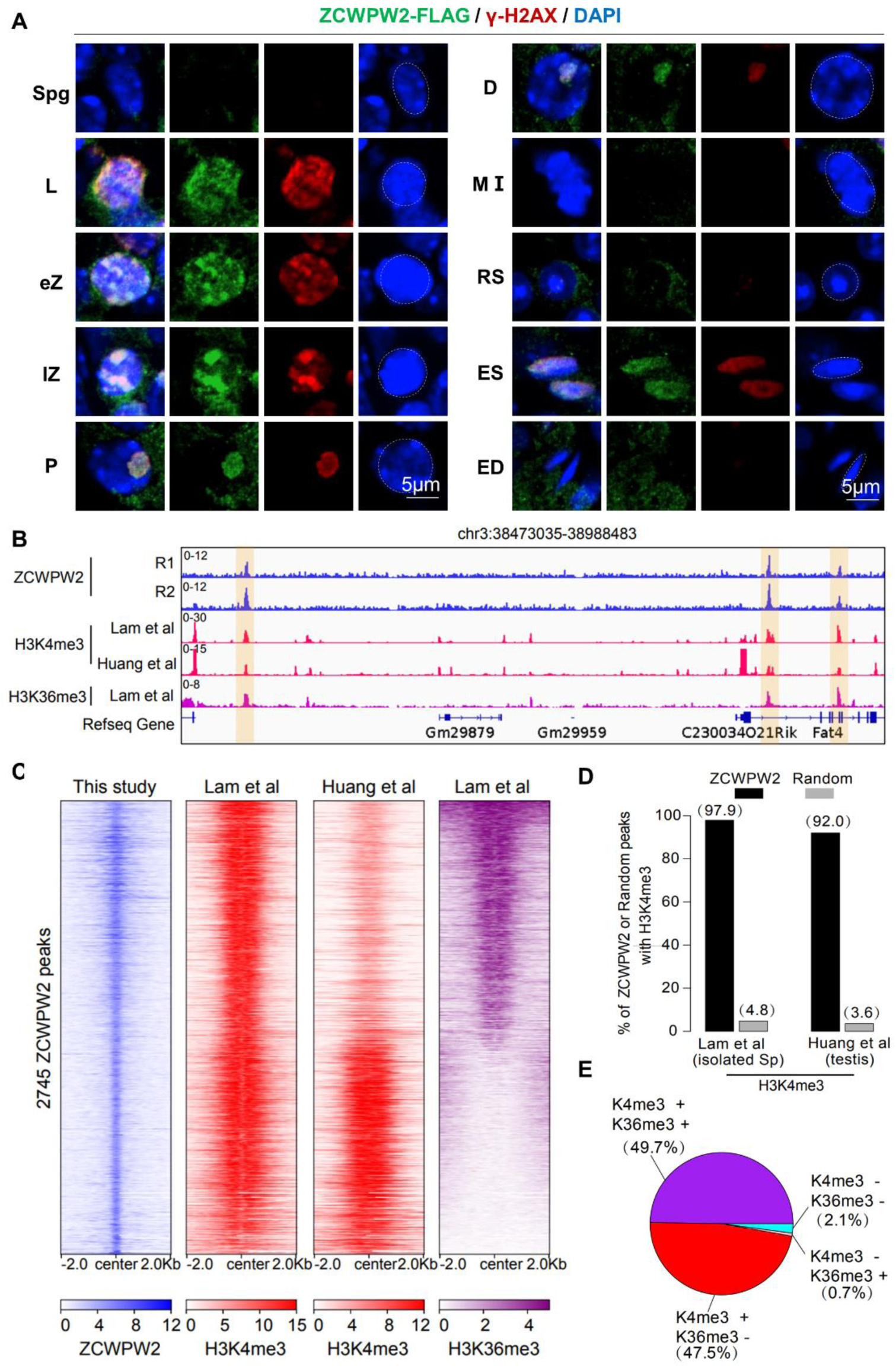
ZCWPW2 binds to sites with both H3K4me3 and H3K36me3 marks. **A.** Double immunostaining for FLAG and ãH2AX in *Zcwpw2*-FLAG knock in mouse germ cells from testis sections of the indicated stages, showing ZCWPW2’s dynamic expression profile during spermatogenesis. Nuclei were counterstained with DAPI. Spg, Spermatogonia; L, leptotene; eZ, early zygotene; lZ, late zygotene; P, pachytene; D, diplotene; MⅠ, metaphase Ⅰ; RS, round spermatids; ES, elongating spermatids; ED, elongated spermatids. Scale bar, 5μm. **B.** ChIP-seq genome snapshot of the distribution of ZCWPW2, H3K4me3 and H3K36me3 peaks in wild type C57BL/6 mice. Overlapping regions are indicated by orange shaded areas. R1 and R2 represent two independent replicates. The H3K4me3 and H3K36me3 tract labeled Lam et al. was generated with isolated stage-specific (SCP3^+^H1T^-^) spermatocyte nuclei (Lam et al., 2019). All other datasets were derived from whole testis samples. **C.** Heatmap showing the ZCWPW2, H3K4me3 and H3K36me3 signals on all the detected ZCWPW2 peaks in wild type testes. **D.** Bar plot showing the percentage of ZCWPW2 peaks or the random regions overlapped with H3K4me3 peaks in testis (Huang et al) and isolated stage-specific spermatocyte (Lam et al). **E.** Pie chart showing the ratio of four ZCWPW2 peak groups determined by their overlap with H3K4me3 and H3K36me3 peaks generated with isolated stage-specific spermatocyte nuclei (Lam et al.). The ‘+” indicates overlap, while ‘−” indicates no overlap.

Similar to many chromatin-related proteins, the ZCWPW2 protein is primarily located in the nuclear region, rather than in the cytosol. To confirm the nuclear localization of ZCWPW2, we performed western blot of ZCWPW2 in cytosolic and nuclear fractions from mouse testes, wherein ZCWPW2 was predominantly localized in the nuclear fraction (Fig.S2A). This observation suggests that ZCWPW2 may play a role in nuclear events, such as chromatin activity. To further confirm this observation, we then examined the chromatin-binding activity of ZCWPW2 and several known chromatin-binding proteins by using different washing conditions. As expected, ZCWPW2 exhibited obvious chromatin-binding activity (Fig.S2B). It has been previously discovered that the CW and PWWP domains can bind to H3K4me3 and H3K36me3, respectively[34]. To investigate whether ZCWPW2 protein harboring CW and PWWP domains is capable of binding H3K4me3 and H3K36me3.The immunofluorescence results confirmed that ZCWPW2 was highly colocalized with H3K4me3 and H3K36me3 in the nuclei of spermatocytes (Fig.S2C). This colocalization in meiotic germ cells supports its role as a dual reader of H3K4me3 and H3K36me3 during spermatogenesis.

In order to identify ZCWPW2 binding sites in the genome, we performed ChIP-seq for ZCWPW2 in the mouse testis. In total, we detected 2745 ZCWPW2 peaks (binding sites) (Fig.1B-C & Supplementary Table S1). To define the chromatin context of these sites, we integrated publicly available epigenomic datasets from spermatogenesis, including profiles of histone modifications and chromatin-associated factors (Supplementary Table S2)[35]. Intriguingly, 97.9% and 92.0% of ZCWPW2 peaks are marked with strong H3K4me3 signal in isolated spermatocytes and whole testis, respectively (Fig.1D). Furthermore, 49.7% of ZCWPW2 peaks are marked with both H3K4me3 and H3K36me3 while 47.5% of ZCWPW2 peaks are exclusively marked with H3K4me3 (Fig.1E). Notably, when both histone modifications are present, the binding regions of ZCWPW2 are significantly broader, coupled with an increase in the binding intensity (Fig.S2D). These results suggest that the dual histone methylation reader ZCWPW2 binds to sites with both H3K4me3 and H3K36me3 marks in mouse testes.

### ZCWPW2 localizes to DSB hotspots in a PRDM9-dependent manner

Considering that ZCWPW2 can bind to chromatin, we subsequently performed binding motif analysis for ZCWPW2 peaks. In this analysis, the de novo motif "GGTAGTAKCATC" was identified as the motif that could be most significantly enriched by the ZCWPW2 peaks (Fig.S3A). Unexpectedly, among the top five analogous motifs, we discovered that the ZCWPW2 binding motif bore the greatest similarity to the PRDM9 binding motifs (Fig.S3B). Besides, ZCWPW2 proteins displayed a similar expression pattern to PRDM9 in the testis across different developmental stages (Fig.S1B). These results suggest that there may be some degree of colocalization between ZCWPW2 and PRDM9. To validate this assumption, we compared the overlap between ZCWPW2 peaks and the PRDM9 peaks identified in vivo by ChIP-seq[36] and in vitro by affinity-seq[37]. 32.1% of ZCWPW2 peaks overlapped with the in vivo PRDM9 peaks while 44.6% of ZCWPW2 peaks overlapped with the in vitro PRDM9 peaks (Fig.2A-B). Furthermore, we proceeded to explore the association between PRDM9 and ZCWPW2 in the testis. We categorized PRDM9 peaks into two groups: those with a strong ZCWPW2 signal (Group I) and those with a weak ZCWPW2 signal (Group II) (Fig.2C). It is notable that most of PRDM9 peaks are marked with H3K4me3 and H3K36me3, which is consistent with previous findings that PRDM9 can catalyze the histone methylations. However, both the PRDM9 and histone methylation signals are relatively higher in group I than group II (Fig.2C and Fig.S3C). These results indicated that the genomic sites co-localized by PRDM9 and ZCWPW2 show stronger histone methylation signals.

**Figure 2.**
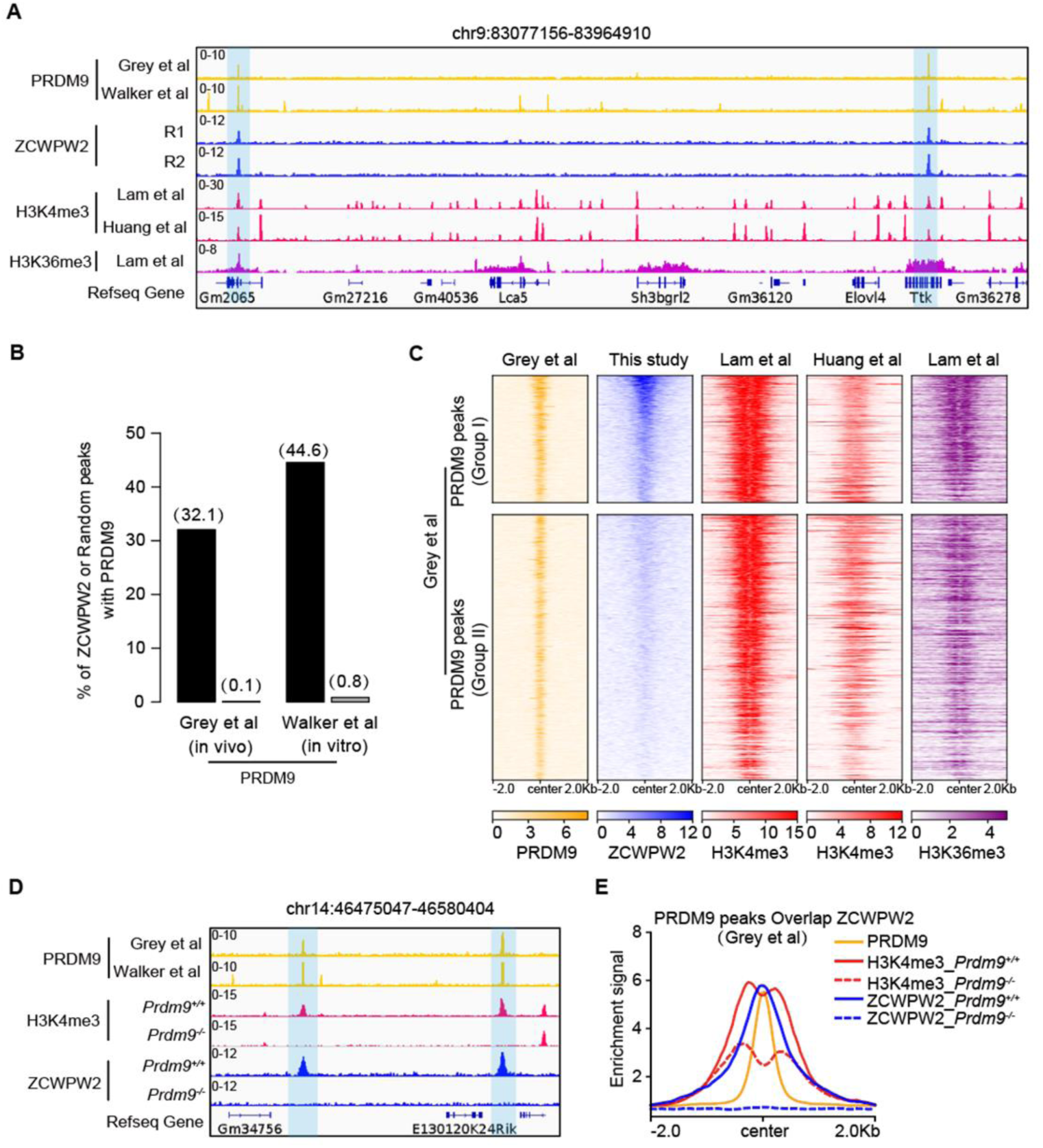
ZCWPW2 localizes to DSB hotspots in a PRDM9-dependent manner. **A.** Genome browser view of PRDM9, ZCWPW2, H3K4me3 and H3K36me3 signals in wild type mouse testes or spermatocytes. The PRDM9 datasets are from in vivo ChIP-seq data generated using an anti-PRDM9 antibody in testis (Grey et al., 2017) and in vitro affinity-seq data (Walker et al., 2015). The blue shadows indicate the genomic regions with PRDM9, ZCWPW2, H3K4me3 and H3K36me3 peaks. **B.** Bar plot showing the percentage of ZCWPW2 peaks or the random regions overlapped with PRDM9 peaks in vivo (Grey et al) and in vitro (Walker et al). **C.** Heatmap showing the PRDM9, ZCWPW2, H3K4me3 and H3K36me3 signals on all the PRDM9 peaks in wild type testes. The PRDM9 peaks are classified into two groups including overlap with ZCWPW2 peaks (Group I) and without ZCWPW2 peaks (Group II). **D.** Genome browser view of H3K4me3 and ZCWPW2 signal on PRDM9 binding regions in wild type and *Prdm9^-/-^*testis. The blue shadows indicate the genomic regions with ZCWPW2 and H3K4me3 peaks in wild type testis, but lost in *Prdm9^-/-^* testis. **E.** Profile plot comparing H3K4me3 and ZCWPW2 signal on the Group I PRDM9 peaks between wild type and *Prdm9^-/-^* testis.

As the aforementioned observation that PRDM9 can catalyze histone methylations at its binding sites, and that ZCWPW2 can recognize these histone methylations, we then hypothesized whether PRDM9 could regulate the binding of ZCWPW2. To validate this hypothesis, we proceeded with the knockout of *Prdm9* in mice. Western blot analysis with mouse testis confirmed that PRDM9 has been successfully knocked out, with no observable impact on the expression levels of ZCWPW2 (Fig.S3D). After the loss of PRDM9, it was obvious that both the H3K4me3 and ZCWPW2 signals decreased at PRDM9 binding sites (Fig.2D and 2E) [38, 39]. Collectively, ZCWPW2 localizes to DSB hotspots, and the ZCWPW2 binding ability is promoted by the histone modification activity of PRDM9.

### ZCWPW2 is essential for spermatogenesis and male fertility

To further analyze the function of ZCWPW2 during spermatogenesis, we generated a *Zcwpw2*-deficient mouse model (hereafter *Zcwpw2^-/-^*) lacking exon 2 to exon 6 of *Zcwpw2* (Fig.3A). The complete loss of ZCWPW2 protein in mouse testes confirmed the successful generation of the *Zcwpw2* knockout mouse model (Fig.S4A-B). Although *Zcwpw2^-/-^* male mice displayed a normal external phenotypic appearance, they were found to be completely infertile, which was accompanied by a significant reduction in testis size (Fig.3B-E). We subsequently examined the phenotypic alterations in mouse germ cells following the knockout of *Zcwpw2*. Histological analysis showed that spermatogenesis in *Zcwpw2^-/-^* males was impaired compared to *Zcwpw2^+/+^* males (Fig.3F). In *Zcwpw2^-/-^* males, the seminiferous tubules lacked post-meiosis spermatids (Fig.3F, asterisk) and contained apoptotic cells (Fig.3F, arrows) or were almost empty (Fig.3F, arrowhead). There were no spermatozoa in the *Zcwpw2^-/-^* epididymis (Fig.3G), thereby providing compelling evidence that *Zcwpw2* is essential for the process of spermatogenesis in mice.

**Figure 3.**
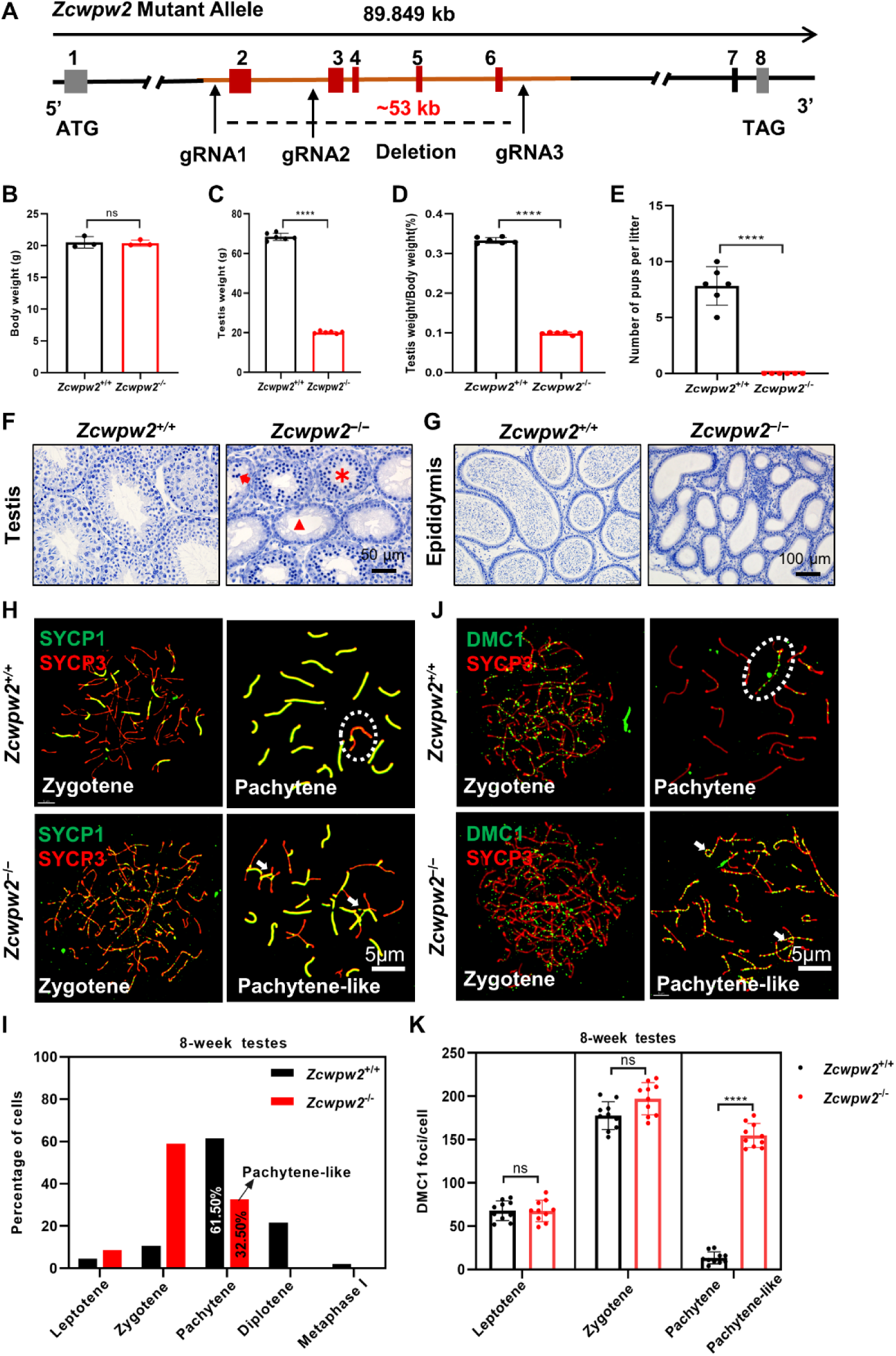
ZCWPW2 is essential for spermatogenesis and male fertility. **A.** Schematic representation of the CRISPR/Cas9 genome editing system for generating *Zcwpw2^−/−^* mice. **B.** The body weights of *Zcwpw2^+/+^* and *Zcwpw2^-/-^* mice. Data presented as mean ± SEM, n = 3. n.s., no statistical significance. **C.** The testis weights of *Zcwpw2^+/+^* and *Zcwpw2^-/-^* mice. Data presented as mean ± SEM, n = 6. ****P < 0.0001 by Student’s t-test. **D.** The ratio of testis weight/body weight in *Zcwpw2^+/+^* and *Zcwpw2^-/-^* mice. Data presented as mean ± SEM, n = 6. ****P < 0.0001 by Student’s t-test. **E.** Fertility assays were performed using male mice (>8 weeks old) paired with wild type females. The average litter size was recorded, and *Zcwpw2^-/-^* males exhibited complete infertility. Data presented as mean ± SEM, n = 6. ****P < 0.0001 by Student’s t-test. **F.** Hematoxylin staining of adult *Zcwpw2^+/+^* and *Zcwpw2^−/−^*testes. Adult *Zcwpw2^−/−^* testes sections showed near complete arrest of spermatogenesis. Arrows, apoptotic spermatocytes; arrowheads, empty seminiferous tubules; asterisks, seminiferous tubules lacking post-meiotic spermatids. Adult mice aged 6–8 weeks, n = 5 per genotype. Scale bar, 50 μm. **G.** Hematoxylin staining of adult *Zcwpw2^+/+^* and *Zcwpw2^−/−^* epididymides. This spermatogenic arrest resulted in empty epididymides in *Zcwpw2^−/−^* mice. Adult mice aged 6–8 weeks, n = 5 per genotype. Scale bar, 100 μm. **H.** Chromosome spreads of spermatocytes from the testes of adult *Zcwpw2^+/+^* and *Zcwpw2^−/−^*males were immunostained for the SC marker proteins SYCP1 (green) and SYCP3 (red). Representative images of pachytene spermatocytes (circle indicates the XY body) in *Zcwpw2^+/+^* mice with completely synapsed chromosomes and pachytene-like spermatocytes in adult *Zcwpw2^−/−^*mice with incompletely synapsed chromosomes (arrows indicate unsynapsed chromosomes). Scale bar, 5 μm. **I.** Frequencies of meiotic stages in adult *Zcwpw2^+/+^* and *Zcwpw2^−/−^* spermatocytes. The numbers marked in the bars represent the percentage of cells at indicated meiotic stages. For each genotype, at least three mice were analyzed. **J.** Chromosome spreads of spermatocytes from the testes of adult *Zcwpw2^+/+^* and *Zcwpw2^−/−^*males were immunostained for DMC1 (green) and SYCP3 (red). Representative images are shown for spermatocytes at the zygotene, pachytene (circle indicates the XY body), and pachytene-like (arrows indicate the DMC1 signal) stages of the two genotypes. All experiments were performed on adult mice (6-8 weeks), n = 3 per genotype. Scale bar, 5 μm. **K.** Each dot represents the number of DMC1 foci per cell, with black dots indicating *Zcwpw2^+/+^* spermatocytes and red dots indicating *Zcwpw2^−/−^* spermatocytes. Solid lines show the mean and SD of the foci number for each group of spermatocytes. P values were calculated by Student’s t test. Adult mice (6-8 weeks), n = 3 per genotype.

TUNEL staining of testicular sections revealed that, starting from PD12, *Zcwpw2^-/-^* mice exhibited a higher number of apoptotic cells in their seminiferous tubules compared to *Zcwpw2^+/+^* mice (Fig.S4C-D). These apoptotic cells were predominantly localized in leptotene and zygotene spermatocytes, suggesting that ZCWPW2 may influence early spermatogenesis. To further validate the early germ cell loss in *Zcwpw2^-/-^* testes, we performed immunofluorescence staining on testicular sections from *Zcwpw2^+/+^*and *Zcwpw2^-/-^* mice at various postnatal ages, using GCNA1 (a germ cell marker) and SOX9 (a Sertoli cell marker). In *Zcwpw2^-/-^*mice aged PD12 to PD16, a significant reduction in germ cells was observed, while the number of Sertoli cells remained unchanged (Fig.S5A-D). Collectively, these findings indicate that ZCWPW2 deficiency results in an early loss of meiotic spermatocytes, which is associated with impaired spermatogenesis and male infertility in *Zcwpw2^-/-^* mice.

### Deficiency of ZCWPW2 disrupts chromosomal synapsis

To explore the underlying cause of spermatogenesis disorders in *Zcwpw2^-/-^*mice, we initially examined the chromosomal status during the prophase I stage of meiosis through immunostaining of synaptonemal complex (SC) proteins. The SC is composed of two axial elements (AEs) and one central element (CE), which are interconnected by transverse filaments[38, 39]. In wild-type spermatocytes, the AEs distinguished by the presence of synaptonemal complex protein 3 (SYCP3) exhibited complete formation at the leptotene stage, concurrent with a gradual progression of chromosomal synapsis, as evidenced by the emergence of CEs along with the SYCP1 signals (Fig.3H). At the pachytene stage, *Zcwpw2^+/+^* spermatocytes achieved complete autosomal synapsis, as demonstrated by the continuous presence of the SYCP1 signal (Fig. 3H). In *Zcwpw2^-/-^* spermatocytes, the formation of AEs also took place during the leptotene stage, and concomitantly, the appearance of SYCP1 on the paired chromosomes was observed, indicating the initiation of synapsis to some extent (Fig.3H). Nonetheless, a significant portion of autosomes in *Zcwpw2^-/-^* spermatocytes remained unsynapsed in comparison to the synapsed chromosomes (Fig.3H). These mutant spermatocytes remained at a stage characterized as "pachytene-like" defined by the presence of more than five synapsed chromosome pairs per cell. We subsequently conducted a quantitative analysis of synapsed chromosome pairs in each nucleus of *Zcwpw2^+/+^* and *Zcwpw2^-/-^*testes at eight weeks of age (Fig.S5E). In the *Zcwpw2^+/+^* testes, a substantial majority of cells (91.3%, 137/150) exhibited complete synapsis across all chromosome pairs. However, none of the 150 spermatocytes examined in the *Zcwpw2^-/-^* testes exhibited full synapsis, and there was merely an average of eight paired synapsed chromosomes per cell (Fig.S5E). Moreover, we extended our analysis to include the proportion of spermatocytes across different meiotic stages in *Zcwpw2^+/+^* and *Zcwpw2^-/-^* testes at postnatal eight weeks. It was observed that spermatocytes in *Zcwpw2^-/-^* testes were unable to progress beyond the pachytene stage, and 32.5% of these spermatocytes were halted at the pachytene-like stage (Fig.3I, pachytene-like). This is in stark contrast to the *Zcwpw2^+/+^* spermatocytes, where 61.5% reached the pachytene stage at postnatal eight weeks (Figure 3I). Collectively, these results suggested that ZCWPW2 deficiency leads to disrupted chromosomal synapsis, thereby impeding spermatogenesis.

### ZCWPW2 deficiency impairs meiotic recombination in spermatocytes

Having demonstrated that ZCWPW2 promotes the completion of synapsis during meiotic prophase I in male mice, we next turned to the assessment of DSB repair. Spermatocyte spreads from the testes of adult *Zcwpw2^+/+^* and *Zcwpw2^-/-^* mice were stained for γH2AX, a marker of DSBs[40]. While DSB formation occurred normally across all genotypes (Fig.S6A), clear differences emerged between pachytene *Zcwpw2^+/+^* spermatocytes and pachytene-like *Zcwpw2^-/-^*spermatocytes. In the *Zcwpw2^+/+^* pachytene spermatocytes, no obvious γH2AX signal was detected on autosomes; however, a γH2AX signal persisted on the sex chromosomes, corresponding to the XY body and indicating sex chromosome silencing. By contrast, both autosomes and sex chromosomes retained pronounced γH2AX signals in *Zcwpw2^-/-^* pachytene-like spermatocytes, with no XY bodies observed (Fig.S6A).

We next performed immunofluorescence staining for phosphorylated ATM (pATM), which marks activated ATM at DNA double-strand break sites[41]. In *Zcwpw2^+/+^* spermatocytes, pATM signals were abundant along chromosome axes at the zygotene stage and were nearly completely absent by the pachytene stage, consistent with the normal temporal dynamics of ATM activation during meiosis. In *Zcwpw2^-/-^* spermatocytes, pATM localization at zygotene was similar to that observed in *Zcwpw2^+/+^*controls. However, spermatocytes from *Zcwpw2^-/-^*mice were arrested at a pachytene-like stage with defective synapsis, and pATM signals persisted abnormally on unsynapsed chromosome axes, failing to be downregulated as seen in *Zcwpw2^+/+^* pachytene cells (Fig.S6B). These findings indicate that loss of ZCWPW2 results in sustained ATM activation in meiotic spermatocytes, which correlates with meiotic arrest at the pachytene-like stage.

To precisely assess DSB repair and recombination, we performed immunostaining for RPA2, RAD51, and DMC1. Initially, the number of RPA2 foci was comparable between *Zcwpw2^+/+^* and *Zcwpw2^-/-^* spermatocytes at the leptotene and zygotene stages (Fig.S6C-D). However, as meiotic recombination progressed, a marked reduction in RPA2 foci was observed in *Zcwpw2^+/+^* spermatocytes at the pachytene stage, whereas these foci persisted in *Zcwpw2^-/-^* spermatocytes (Fig.S6C-D), suggesting inefficient RPA clearance. Similarly, the counts of RAD51 and DMC1 foci were equivalent in both genotypes during leptotene and zygotene stages (Fig.3J-K, S6E-F). At the late pachytene or pachytene-like stages, however, the numbers of RAD51 and DMC1 foci were significantly higher in *Zcwpw2^-/-^* spermatocytes compared to *Zcwpw2^+/+^* spermatocytes (Fig.3J-K, S6E-F). This indicates that *Zcwpw2^-/-^* spermatocytes face obstacles in recombination and DSB repair, as reflected by the sustained presence of RAD51/DMC1 recombinases on their chromosomes.

Seeking to further assess the functional contributions of ZCWPW2 in meiotic recombination, we examined spermatocyte chromosome spreads from adult *Zcwpw2^+/+^* and *Zcwpw2^-/-^* mice via immunostaining for the recombination regulators MSH4, RNF212, and the Holliday junction resolution marker MLH1[42]. Immunostaining of MSH4 and RNF212 revealed that the recombination machinery assembled properly in both *Zcwpw2^+/+^* and *Zcwpw2^-/-^* spermatocytes at leptotene and zygotene stages. While these signals gradually diminished in *Zcwpw2^+/+^*pachytene spermatocytes, they remained abnormally sustained on chromosomes of pachytene-like *Zcwpw2^-/-^*spermatocytes (Fig.S7A-D). During the pachytene stage, *Zcwpw2^+/+^*spermatocytes displayed prominent MutL homolog 1 (MLH1) foci, signifying the sites of crossovers (Fig.S7E-F). In contrast, *Zcwpw2^-/-^* spermatocytes were unable to progress to this stage, leading to the absence of crossovers and consequently, the non-appearance of MLH1 foci (Fig.S7E-F). Taken together, these findings suggest that the deficiency in ZCWPW2 may result in impaired DSB repair and recombination, consequently hindering the progression of meiosis.

### Either H3K4me3 or H3K36me3 reader function of ZCWPW2 is essential for meiotic recombination

Having thus established that ZCWPW2 binds to sites marked by both H3K4me3 and H3K36me3 in a PRDM9-dependent manner, and that loss of *Zcwpw2* in male mice led to a complete failure of synapsis, this failure caused meiotic arrest at the "pachytene-like" stage, accompanied by incomplete DSB repair and absence of crossover formation, ultimately resulting in male infertility. In light of the known ability of ZCWPW2 to recognize epigenetic methylation modification marks, and to investigate the reader function of ZCWPW2, we first examined its sequence conservation across diverse species. The results demonstrated that the ZCWPW2 amino acid sequence is highly conserved, particularly within the CW and PWWP domains (Fig.S8A). Alignment of these domains among mouse paralogs further identified a set of highly conserved residues (Fig.S8B-C). Guided by the available human crystal structures of the CW and PWWP domains, we employed Alpha Fold 3 to generate structural models for the corresponding mouse ZCWPW2 domains. Structural analysis revealed that residues W15, E61, and F63 in the CW domain, as well as W94, W97, and F126 in the PWWP domain, occupy key positions within the binding pockets (Fig.S9A). Based on these observations, we designed a knock-in strategy to generate a H3K4me3 reader-dead ZCWPW2 mutant mouse line (hereafter *Zcwpw2^CW-KI^*) and a H3K36me3 reader-dead ZCWPW2 mutant mouse line (hereafter *Zcwpw2^PWWP-KI^*), respectively. Specifically, *Zcwpw2^CW-KI^* mutant had three mutations W15I/E61R/F63P and *Zcwpw2^PWWP-KI^* mutant had three mutations W94I/W97A/F126A (Fig.S9B-C).

Western blot analysis confirmed the absence of the ZCWPW2 protein in *Zcwpw2^-/-^* testes, while the ZCWPW2 ^W15I/E61R/F63P^ or ZCWPW2 ^W94I/W97A/F126A^ variant protein were expressed at a level similar to that of the wild type protein (Fig.S4A and S10A). Consistent with the western blot data, immunofluorescence staining of fixation sections from 8-week-old *Zcwpw2^+/+^*, *Zcwpw2^-/-^*, *Zcwpw2^CW-KI^*and *Zcwpw2^PWWP-KI^* mouse testes revealed that the ZCWPW2 protein was undetectable in *Zcwpw2^-/-^* spermatocytes but could still be found in *Zcwpw2^CW-KI^* and *Zcwpw2^PWWP-KI^* mutant spermatocytes (Fig.S4B and S10B). After confirming that the ZCWPW2 mutant protein could be expressed normally in *Zcwpw2^CW-KI^*and *Zcwpw2^PWWP-KI^* mice, we prepared testis sections from 8-week-old *Zcwpw2^+/+^*, *Zcwpw2^-/-^* and the new *Zcwpw2^CW-KI^* and *Zcwpw2^PWWP-KI^* mice line. Hematoxylin staining showed that spermatogenesis was disrupted in both the *Zcwpw2^CW-KI^* and *Zcwpw2^PWWP-KI^* mice. Compared with the *Zcwpw2^+/+^*mice, the seminiferous tubules of the *Zcwpw2^CW-KI^* and *Zcwpw2^PWWP-KI^*mice lacked post-meiotic cell types, contained apoptotic cells, or were nearly empty. Furthermore, the *Zcwpw2^+/+^* epididymis was full of sperm, but there were no obvious sperm detected in either the *Zcwpw2^CW-KI^* or *Zcwpw2^PWWP-KI^* samples, suggesting meiotic arrest in these reader-dead mutant mice (Fig. 4A). Consistent with the phenotypes observed in *Zcwpw2^-/-^* mice, immunostaining of SYCP1/SCYP3 in *Zcwpw2^CW-KI^*or *Zcwpw2^PWWP-KI^* spermatocytes revealed identical defects in chromosomal synapsis (Fig. 4B). While *Zcwpw2^+/+^* spermatocytes achieved complete autosomal synapsis at the pachytene stage, both knock-in mutant lines exhibited only partial synapsis and were arrested at a pachytene-like stage, fully recapitulating the synapsis failure seen in *Zcwpw2^-/-^* mice. Consistent with this synapsis defect, immunostaining of γ-H2AX and pATM in *Zcwpw2^CW-KI^*or *Zcwpw2^PWWP-KI^* spermatocytes revealed identical defects in DSB repair (Fig.S11A-B). While γ-H2AX and pATM signals were normally restricted to the XY body in *Zcwpw2^+/+^* pachytene spermatocytes, both markers persisted abnormally on unsynapsed chromosome axes in pachytene-like arrested *Zcwpw2^CW-KI^* or *Zcwpw2^PWWP-KI^*spermatocytes, mirroring the persistent DNA damage response seen in *Zcwpw2^-/-^*mice (Fig.S11A-B).

**Figure 4.**
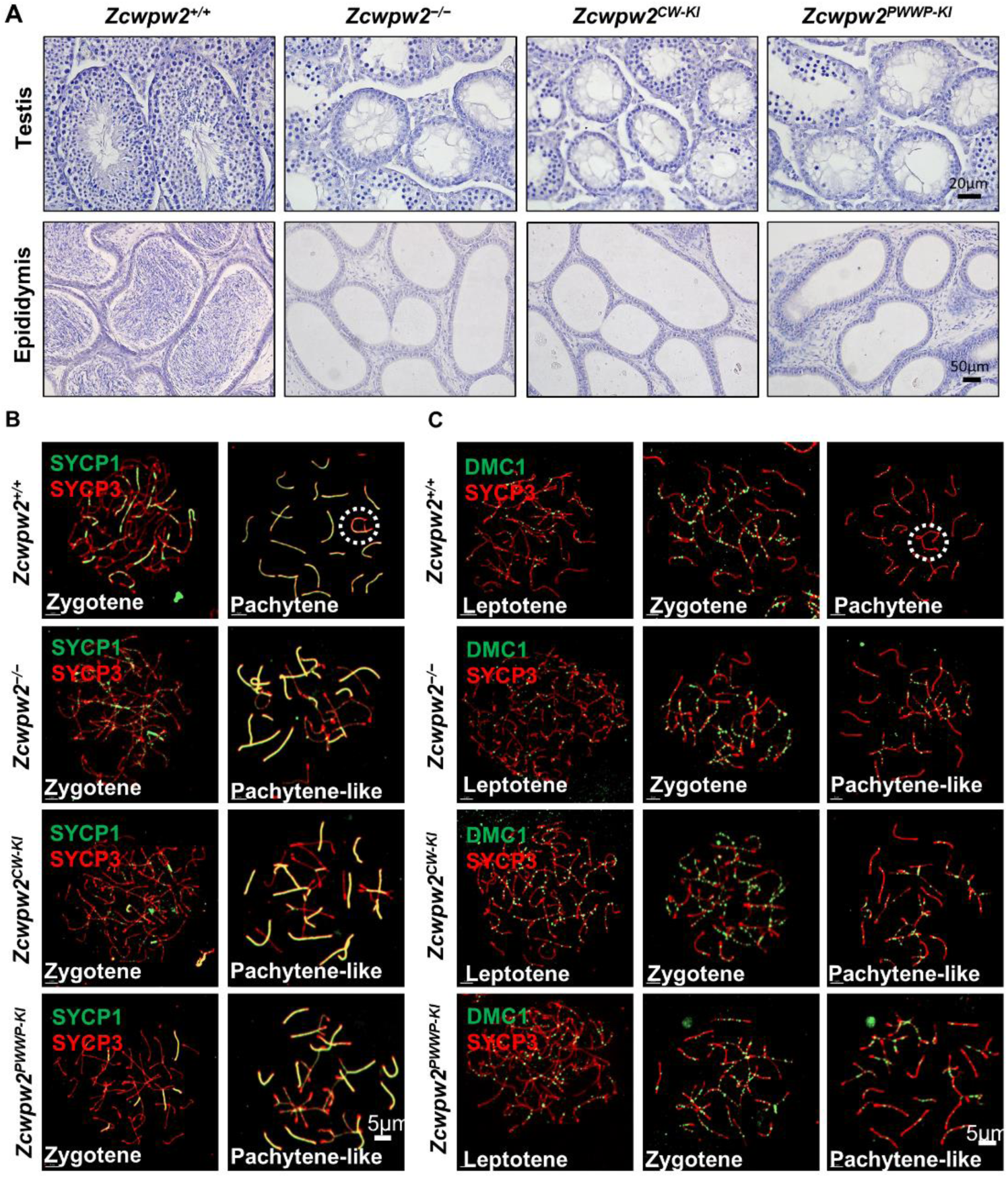
Either H3K4me3 or H3K36me3 reader function of ZCWPW2 is essential for meiotic recombination. **A.** Hematoxylin staining of testes and epididymides from adult *Zcwpw2^+/+^*, *Zcwpw2^−/−^*, *Zcwpw2^CW-KI^*, and *Zcwpw2^PWWP-KI^* mice. *Zcwpw2^CW-KI^* and *Zcwpw2^PWWP-KI^*mice exhibited a similar spermatogenic phenotype to *Zcwpw2^−/−^*mice, with near-complete arrest of spermatogenesis in testicular sections. Scale bar, 20 μm for testes and 50 μm for epididymides. **B.** Chromosome spreads of spermatocytes from the testes of adult *Zcwpw2^+/+^*, *Zcwpw2^−/−^*, *Zcwpw2^CW-KI^*, and *Zcwpw2^PWWP-KI^*males were immunostained for the SC marker proteins SYCP1 (green) and SYCP3 (red). Representative images are shown for spermatocytes at the leptotene, zygotene, pachytene (circle indicates the XY body), and pachytene-like stages across the four genotypes. Scale bar, 5μm. **C.** Chromosome spreads of spermatocytes from the testes of adult *Zcwpw2^+/+^*, *Zcwpw2^−/−^*, *Zcwpw2^CW-KI^*, and *Zcwpw2^PWWP-KI^* males were immunostained for DMC1 (green) and SYCP3 (red). Representative images are shown for spermatocytes at the leptotene, zygotene, pachytene (circle indicates the XY body), and pachytene-like stages across the four genotypes. Scale bar, 5μm.

We next analyzed the dynamics of meiotic recombination factors in reader-dead mutant mice, which fully recapitulated the *Zcwpw2^-/-^* phenotype (Fig.4C and Fig.S12A-B). For RPA2, RAD51, and DMC1, initial loading at leptotene and zygotene stages was comparable between wild-type and reader-dead mutant spermatocytes; however, these foci remained abnormally elevated on unsynapsed chromosome axes in pachytene-like *Zcwpw2^CW-KI^*or *Zcwpw2^PWWP-KI^* spermatocytes, indicating impaired DSB repair and unresolved recombination intermediates, identical to that observed in *Zcwpw2^-/-^* mice (Fig.4C and Fig.S12A-B). MSH4 and RNF212 assembled normally at zygotene in both wild-type and reader-dead mutant spermatocytes but failed to be downregulated in pachytene-like mutant spermatocytes, in contrast to their gradual disappearance in wild-type pachytene cells (Fig.S13A-B). MLH1 foci—markers of mature crossovers—were completely absent in both reader-dead mutant mice, as reader-dead mutant spermatocytes were arrested at a pachytene-like stage and failed to progress to the crossover formation stage, fully recapitulating the *Zcwpw2^-/-^*phenotype (Fig.S14A). Collectively, disruption of either the CW or PWWP domain of ZCWPW2 leads to severe defects in meiotic chromosome synapsis and DSB repair, impairs meiotic recombination, and fully recapitulates the complete spermatogenic failure phenotype observed in *Zcwpw2^-/-^*mice. Thus, either H3K4me3 or H3K36me3 reader function of ZCWPW2 is essential for meiotic recombination.

### ZCWPW2 is required for normal DSB formation at meiotic hotspots

In mammals, the catalytic formation of H3K4me3 and H3K36me3 by PRDM9 (writer) determines the formation of DSBs at meiotic recombination hotspots[28]. Nevertheless, the precise involvement of histone methylations in DSB formation remain largely unexplored. Our data reveal that ZCWPW2 could bind to the meiotic hotspots through recognizing PRDM9-deposited histone methylations. Thus, we sought to explore whether ZCWPW2 potentially exerts an influence on the formation of DSBs at meiotic hotspots. Firstly, we measured the overlap between ZCWPW2 peaks and DSB sites which could be represented by the DMC1 peaks or End-seq peaks[36, 43, 44]. About 50% of ZCWPW2 peaks localized at meiotic DSB sites in *Zcwpw2^+/+^* mouse testis (Fig.5A and Fig.S15A). The END-seq signal at DSB sites became stronger along with the increase of ZCWPW2 signal (Fig.5B). These results suggest there may be somewhat putative association between the ZCWPW2 and DSB formation.

**Figure 5.**
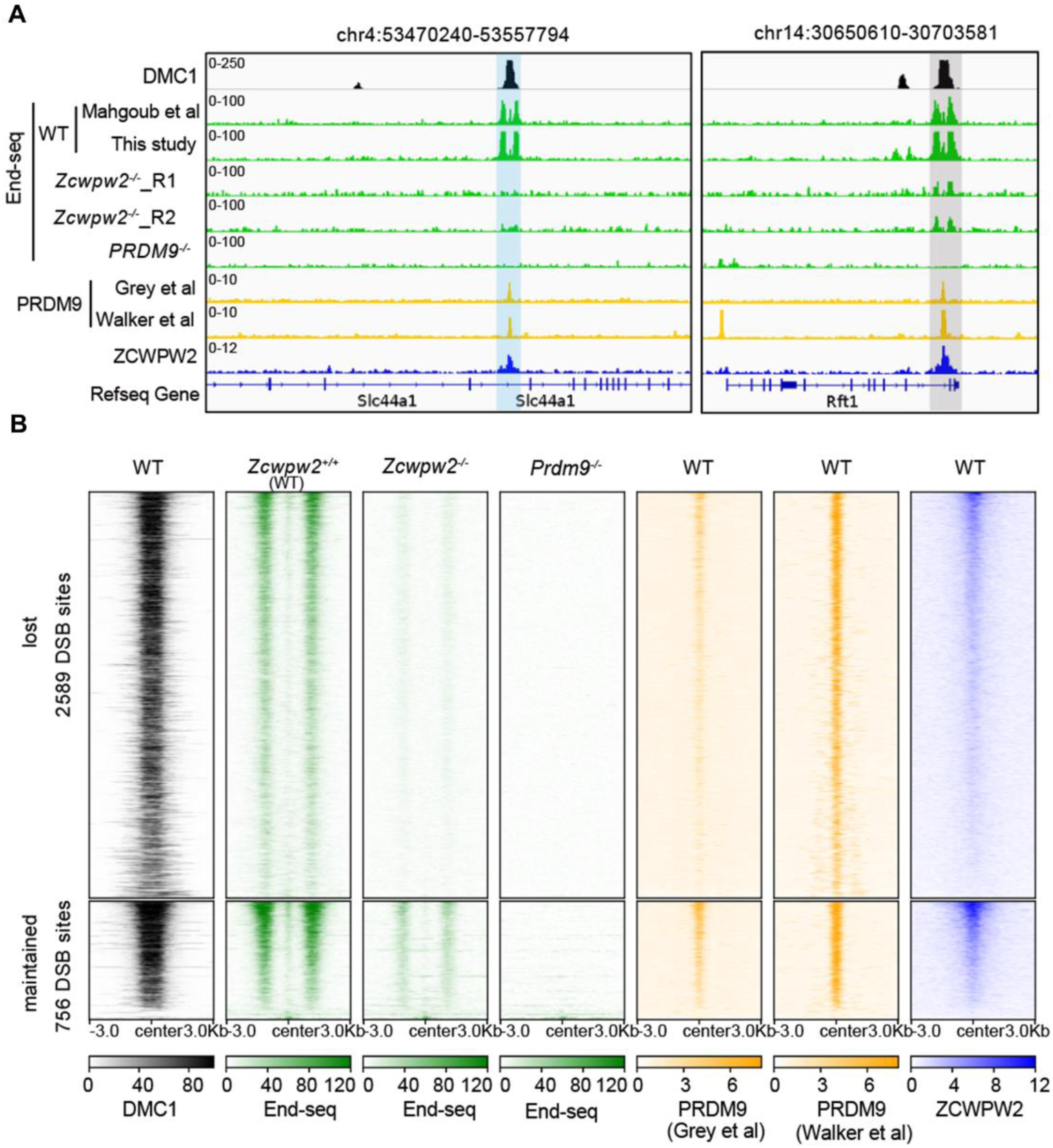
ZCWPW2 is required for normal DSB formation at meiotic hotspots. **A.** Genome browser view of DMC1, PRDM9 and ZCWPW2 signal in wild type testes, Enq-seq signal in wild type, *Zcwpw2^-/-^* and *Prdm9^-/-^* testis. The blue shadows indicate the ZCWPW2 and PRDM9 binding regions with DMC1 and End-seq peaks in wild type testes, but lost in *Zcwpw2^-/-^*and *Prdm9^-/-^* testis. The grey shadows indicate the ZCWPW2 and PRDM9 binding regions with End-seq peaks in wild type and *Zcwpw2^-/-^* testes. The DMC1 and End-seq in wild type testes peaks represent the DSB hotspots. **B.** Heatmap showing the difference of DSB signal on physiological END-seq peaks in *Zcwpw2^-/-^*and *Prdm9^-/-^* testis. The END-seq peaks include those lost and maintained in *Prdm9^-/-^* testis. DMC1, PRDM9, and ZCWPW2 signals in wild-type testes are also shown.

Compared to 3345 END-seq peaks in *Zcwpw2^+/+^* mouse testis, 4966 END-seq peaks could also be identified in *Zcwpw2^-/-^*mouse testis (Fig.S15B). However, among the END-seq peaks in *Zcwpw2^+/+^* mouse testis, 77.4% (2589/3345) END-seq peaks (lost group) could not be detected in *Zcwpw2^-/-^* mouse testis (Fig.5A-B and Fig.S15B). The remaining 22.6% (756/3345) END-seq peaks (maintained group) are still detectable but with a remarkable decrease in the *Zcwpw2^-/-^*mouse testis (Fig.5A-B and Fig.S15B). These results indicate loss of *Zcwpw2* could lead to the failure of DSB positioning at meiotic recombination hotspots. Consistent with a previous report, we also found that all the DSB sites in the lost group exhibit no DSB signal in *Prdm9^-/-^*mouse testis (Fig.5A-B). However, most of the DSB sites in the maintained group also exhibit no DSB signal in *Prdm9^-/-^* mouse testis, rather than relatively weak DSB signal (Fig.5A-B). Collectively, our data reveal that the epigenetic writer-reader axis PRDM9-ZCWPW2 is essential for DSB formation at meiotic recombination hotspots.

Beyond the lost and maintained groups, we detected 4214 new END-seq peaks (gained group) in *Zcwpw2^-/-^* mouse testis (Fig.S15B-C). Almost all of the newly identified END-seq peaks in the gained group exhibit neither DMC1 nor END-seq signals in the *Zcwpw2^+/+^* mouse testis (Fig.S15C-D). These findings suggest that the loss of *Zcwpw2* cause DSBs to occur at incorrect locations, instead of at the meiotic hotspots. Moreover, the DSB sites within the gained group also display DSB signal in *Prdm9^-/-^* mouse testis (Fig.S15C-D), which is consistent with a previous report that loss of *Prdm9* could cause DSBs to occur at incorrect locations[27]. Upon further examination of the genomic distribution of DSB sites across the three groups, we found that the majority of the gained DSB sites are predominantly situated in the promoter regions, unlike the lost and maintained DSB sites (Fig.S15E). Taken together, our findings suggest that the disruption of the epigenetic writer-reader axis, PRDM9-ZCWPW2, can misdirect positioning of DSBs to promoter regions.

### ZCWPW2 physically interacts with multiple DSB machinery complex subunits

Meiotic DSB formation is a tightly regulated process dependent on the coordinated function of axis-associated proteins and the SPO11 topoisomerase complex[7, 8]. As characterized in prior studies, HORMAD1 localizes to meiotic chromosome axes to enforce the fidelity of homologous synapsis and execute key checkpoint control functions; IHO1 serves as a critical adaptor that directly links HORMAD1 to the DSB formation machinery, thereby facilitating efficient SPO11-mediated DSB induction[45, 46]. MEI4 and REC114, as conserved and indispensable core components of the SPO11-activating complex, are essential for the initiation of meiotic recombination and the proper spatial-temporal progression of downstream homologous recombination events[47]. To elucidate the molecular regulatory relationship between ZCWPW2 and this core axis-DSB regulatory network, we performed co-immunoprecipitation (Co-IP) assays in human embryonic kidney 293T (HEK293T) cells combined with yeast two-hybrid (Y2H) analyses to verify their physical interactions.

Co-IP experiments in HEK293T cells demonstrated specific physical associations between ZCWPW2 and the core meiotic recombination factors: immunoprecipitation of HORMAD1 efficiently pulled down ZCWPW2, and conversely, immunoprecipitation of ZCWPW2 also recovered HORMAD1 in the immunocomplex, confirming their direct interaction in mammalian cellular contexts (Fig.6A). We further verified this interaction endogenously under physiological conditions, as Co-IP with specific antibodies demonstrated robust co-precipitation of endogenous ZCWPW2 and HORMAD1 (Fig.S16A). Consistent with the established role of IHO1 as a HORMAD1-interacting regulator of DSB formation, Co-IP assays targeting FLAG-tagged IHO1 revealed robust co-precipitation of ZCWPW2, indicating that ZCWPW2 is stably incorporated into the HORMAD1-IHO1 axis-associated complex (Fig. 6B). Furthermore, Co-IP analyses for FLAG-tagged MEI4 and REC114—core components of the SPO11-dependent DSB formation machinery—showed that ZCWPW2 was specifically co-purified with both MEI4 and REC114 (Fig.6C-D), demonstrating that ZCWPW2 physically associates with the functional meiotic DSB-inducing machinery.

**Figure 6.**
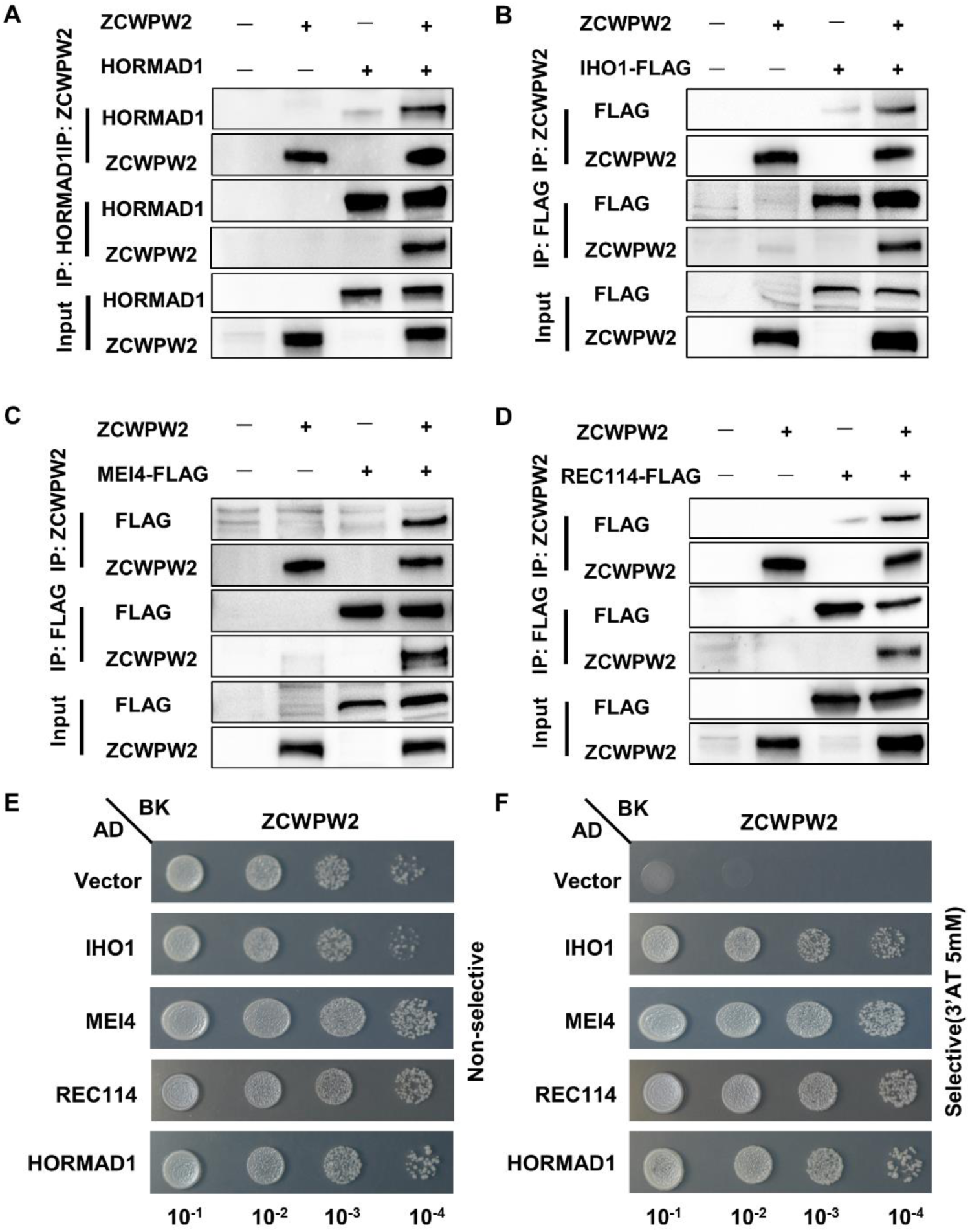
ZCWPW2 physically interacts with multiple DSB machinery complex subunits. **A-D.** Co-IP analysis of the interactions between ZCWPW2 and HORMAD1, IHO1, MEI4 and REC114 in HEK293T cells overexpressing ZCWPW2 and indicated proteins for 48 h. Data are representative of three independent experiments. **E-F.** Yeast two-hybrid analysis. ZCWPW2 was used as bait (BK fusion), while HORMAD1, IHO1, MEI4, and REC114 were used as prey (AD fusion). Yeast transformants were serially diluted (10⁻¹ to 10⁻⁴) and spotted on non-selective and selective (3’AT 5mM) media. Data are representative of three independent experiments.

To independently validate these interactions in a heterologous system, we conducted Y2H assays. Yeast cells co-expressing ZCWPW2 as the bait construct and HORMAD1, IHO1, MEI4, or REC114 as the prey constructs grew robustly on both non-selective (Fig.6E) and stringent selective media (Fig.6F), with growth kinetics comparable to those of positive control groups, whereas empty vector controls exhibited no detectable growth (Fig.6E-F). Collectively, these data establish that ZCWPW2 physically interacts with the entire evolutionarily conserved axis-associated DSB regulatory module—comprising HORMAD1, IHO1, MEI4, and REC114—in heterologous mammalian cell systems, thereby positioning ZCWPW2 as a novel component of the molecular network that governs meiotic DSB formation and the regulation of chromosome synapsis.

## Discussion

Meiotic recombination ensures faithful transmission of the genome by facilitating the pairing, synapsis, and segregation of homologous chromosomes, while also enhancing genetic diversity through the reshuffling of linked alleles[1, 3]. At the molecular level, meiotic recombination begins with programmed DSBs generated during leptotene, which preferentially occur at discrete genomic regions known as hotspots rather than being randomly distributed[48]. In mammals, multiple studies have identified PRDM9 as the key factor defining hotspot identity, through its sequence-specific DNA binding and its enzymatic deposition of H3K4me3 and H3K36me3 at future recombination sites[26]. Nevertheless, how these PRDM9-dependent histone methylation marks are subsequently interpreted to guide physiological DSB formation remains a central unresolved question in the field.

In the present study, we identify ZCWPW2 as a dual histone methylation reader that acts downstream of PRDM9 during meiotic recombination. Our data support a working model in which PRDM9 binds specific DNA motifs and deposits H3K4me3 and H3K36me3 through its PR/SET domain, thereby creating a chromatin environment that marks future meiotic hotspots. ZCWPW2 is then recruited to these sites through its ability to recognize both histone modifications, and contributes to the proper execution of DSB formation and recombination progression. Consistent with this model, ZCWPW2 preferentially occupies PRDM9-dependent hotspot regions, its chromatin association is strongly dependent on PRDM9 function, and its loss leads to defective synapsis, persistent recombination intermediates, failure of crossover formation, and meiotic arrest. These findings place ZCWPW2 as a critical effector that links epigenetic hotspot marking to the downstream meiotic recombination machinery.

A major conceptual implication of our findings is that ZCWPW2 may function, at least in part, as a mammalian counterpart of chromatin–axis coupling factors that connect hotspot-associated histone modifications to the meiotic DSB machinery. In budding yeast, Spp1 recognizes H3K4me3-marked promoter regions and links them to the chromosome axis through interaction with Mer2, thereby facilitating DSB formation at appropriate genomic sites[24, 25]. In mammals, by contrast, hotspot usage has been evolutionarily redirected away from constitutive promoter-associated H3K4me3 sites toward PRDM9-defined loci, indicating that the ancestral H3K4me3-based mechanism has been substantially remodeled[32, 33]. Our data suggest that ZCWPW2 has emerged as a key component of this remodeled system: rather than acting at promoter-associated H3K4me3 sites, ZCWPW2 preferentially reads PRDM9-deposited H3K4me3 and H3K36me3 at meiotic hotspots and functionally connects these chromatin features to axis-associated DSB factors. This model provides a conceptual framework for how mammals may have adapted a conserved chromatin-reading principle to a PRDM9-dependent hotspot specification system.

Our results also refine the mechanistic understanding of how PRDM9-dependent histone methylation contributes to DSB positioning. Although PRDM9 has long been recognized as the writer that establishes hotspot identity, the mechanistic path from histone methylation deposition to SPO11-dependent DNA cleavage has remained unclear. Here, the marked loss of canonical hotspot-associated END-seq signals in *Zcwpw2*-deficient testes, together with the emergence of ectopic promoter-proximal DSBs, indicates that histone methylation marks alone are insufficient to ensure correct DSB placement. Instead, these epigenetic modifications appear to require interpretation by a dedicated reader, namely ZCWPW2, to couple PRDM9-marked chromatin to a DSB-permissive chromosomal axis environment. In this regard, ZCWPW2 should not be viewed merely as a passive binder of histone marks, but rather as an active effector that translates hotspot chromatin identity into spatially accurate meiotic break formation. The physical association of ZCWPW2 with HORMAD1, IHO1, MEI4, and REC114 further supports such a role and places ZCWPW2 within the molecular interface between hotspot chromatin and the conserved DSB machinery.

Another important aspect of this study is that both histone-reading modules of ZCWPW2 are required for its meiotic function. Disruption of either the CW domain or the PWWP domain fully phenocopied the null mutation, leading to defective synapsis, persistent DSB repair defects, loss of crossover formation, and spermatogenic failure. These findings argue that combinatorial recognition of H3K4me3 and H3K36me3 is not merely auxiliary but is essential for ZCWPW2 activity in vivo. Such dual recognition may provide an additional layer of specificity and robustness, ensuring that ZCWPW2 is recruited selectively to authentic PRDM9-defined recombination hotspots rather than to other genomic regions carrying only one of these marks. More broadly, this observation supports the idea that combinatorial histone readout is a central design principle in meiotic chromatin regulation, allowing hotspot-associated epigenetic information to be interpreted with high fidelity.

Several important questions remain to be addressed. Although our genetic, genomic, and biochemical data strongly support an essential role for ZCWPW2 downstream of PRDM9, the precise molecular sequence through which ZCWPW2 coordinates hotspot chromatin, chromosome axes, and the DSB machinery remains to be fully defined. It will be important in future studies to determine whether ZCWPW2 directly stabilizes recruitment of the HORMAD1–IHO1–MEI4–REC114 module to PRDM9-defined hotspots, whether it promotes higher-order chromatin–axis juxtaposition before DSB formation, or whether it contributes to both processes. It will also be important to determine how ZCWPW2 function is coordinated with other PRDM9-associated factors and whether additional histone readers participate in parallel or sequential steps of hotspot activation. More broadly, because meiotic recombination failure is a major cause of infertility, our findings may have implications for reproductive biology and human disease. Defining whether dysfunction of the PRDM9–ZCWPW2 axis contributes to unexplained meiotic arrest or idiopathic male infertility will therefore be an important direction for future investigation.

In summary, our study identifies ZCWPW2 as a dual H3K4me3/H3K36me3 reader that is essential for PRDM9-dependent meiotic DSB formation and recombination progression in mammals. By integrating histone methylation recognition with the axis-associated DSB machinery, ZCWPW2 provides a mechanistic link between hotspot chromatin identity and accurate meiotic break formation. These findings establish ZCWPW2 as a critical downstream effector of PRDM9 and offer new insight into how epigenetic information is interpreted to control the initiation of meiotic recombination.

## Acknowledgements

We would like to thank Wei Wu and Minghan Tong (University of Chinese Academy of Sciences), Jianqiang Bao (University of Science and Technology of China) for technical surport and helpful discussions. We also thank all our colleagues in the Chen laboratory for helpful discussions. We appreciate the support of the Translational Medicine Core Facility of Shandong University for consultation and instrument use.

## Author contributions

T.H. and S.L.Y. conceived and designed the entire project. T.H., S.L.Y., S.Y.W., Z.Y.B. and Z.Q.W. performed most of the experiments, data collection and analysis. K.S.G., M.R.F., and C.Q.H. participated in animal husbandry, genotyping and functional experiments. Y.L.Y., Y.J.S., and X.F. participated in technical guidance and provided advice. H.Z., S.G.Z. and H.B.L. facilitated the study design. T.H., S.L.Y., Z.Q.W., S.Y.W. and Z.Y.B analyzed the data and wrote the manuscript with the assistance of the other authors. Z.-J.C., H.B.L. and T.H. supervised the project. All authors reviewed and approved the final manuscript.

## Data availability

The ZCWPW2 ChIP-seq and End-seq data generated in this study have been deposited in the Genome Sequence Archive (GSA) and are accessible under accession number CRA041943 (https://ngdc.cncb.ac.cn/gsa/s/BSr176bQ). Additionally, all publicly available data utilized in this study are summarized in Supplementary Table S2.

## Supplementary Figures

**Supplementary Figure S1.**
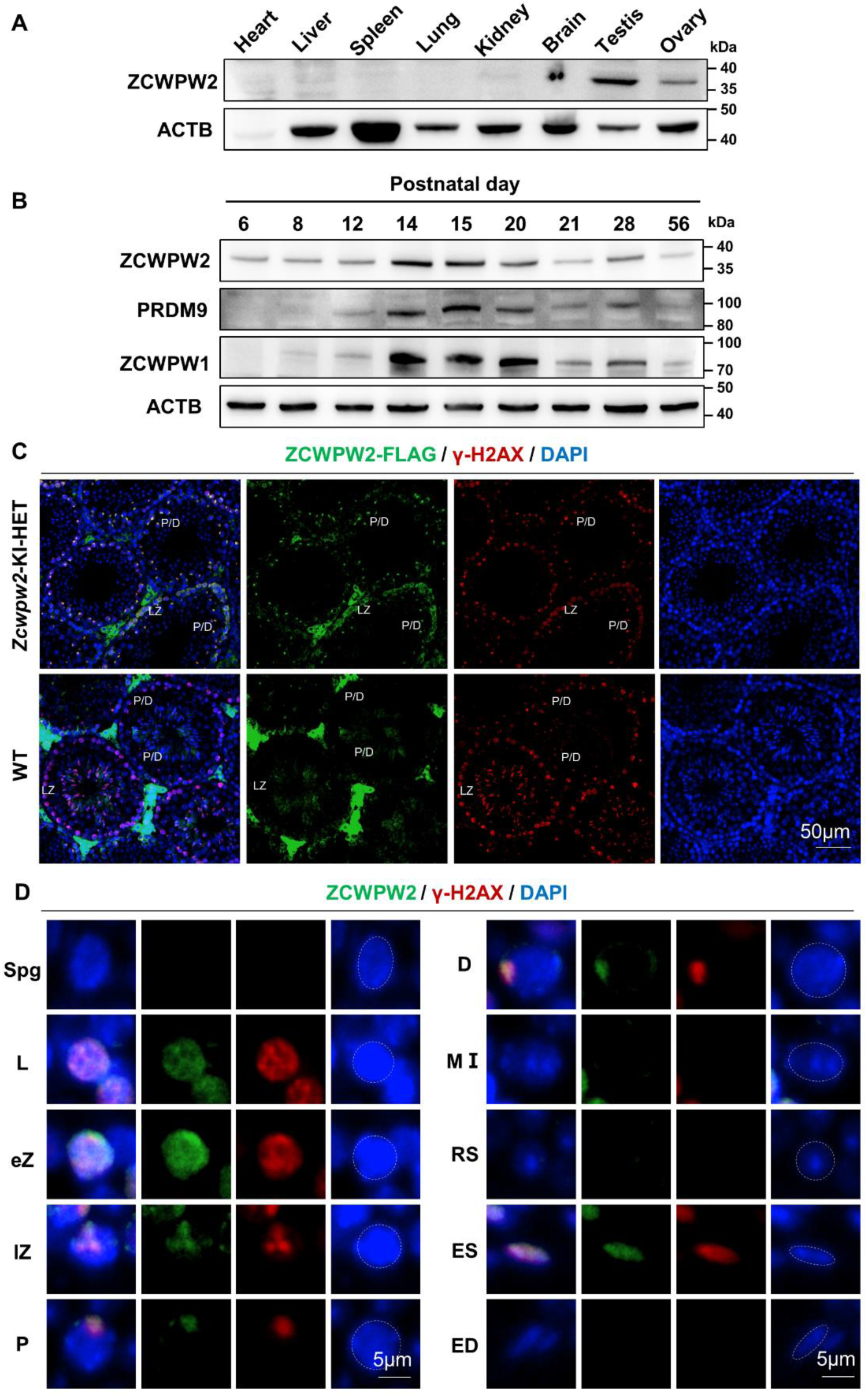
The expression and localization of ZCWPW2 in mice. **A.** Western blotting of ZCWPW2 in different mouse tissues, demonstrating that ZCWPW2 is highly expressed in the testes. **B.** Dynamic expression of ZCWPW2, PRDM9 and ZCWPW1 proteins during spermatogenesis. ZCWPW2 expression increased dramatically at PD14. β-Actin was used as a loading control. **C.** Testis sections from wild type, *Zcwpw2*-FLAG heterozygous knock-in mice were stained with anti-FLAG (green) and γH2AX (red) antibodies. Nuclei were counterstained with DAPI. L/Z, leptotene/zygotene; P/D, pachytene/diplotene. Scale bar, 50 μm. **D.** Double immunostaining against ZCWPW2 and γH2AX in wild type germ cells from testis sections of indicated stages, showing ZCWPW2’s dynamic expression profile during adult spermatogenesis. Nuclei were counterstained with DAPI. Spg, Spermatogonia; L, leptotene; eZ, early zygotene; lZ, late zygotene; P, pachytene; D, diplotene; MⅠ, metaphase Ⅰ; RS, round spermatids; ES, elongating spermatids; ED, elongated spermatids. Scale bar, 5 μm.

**Supplementary Figure S2.**
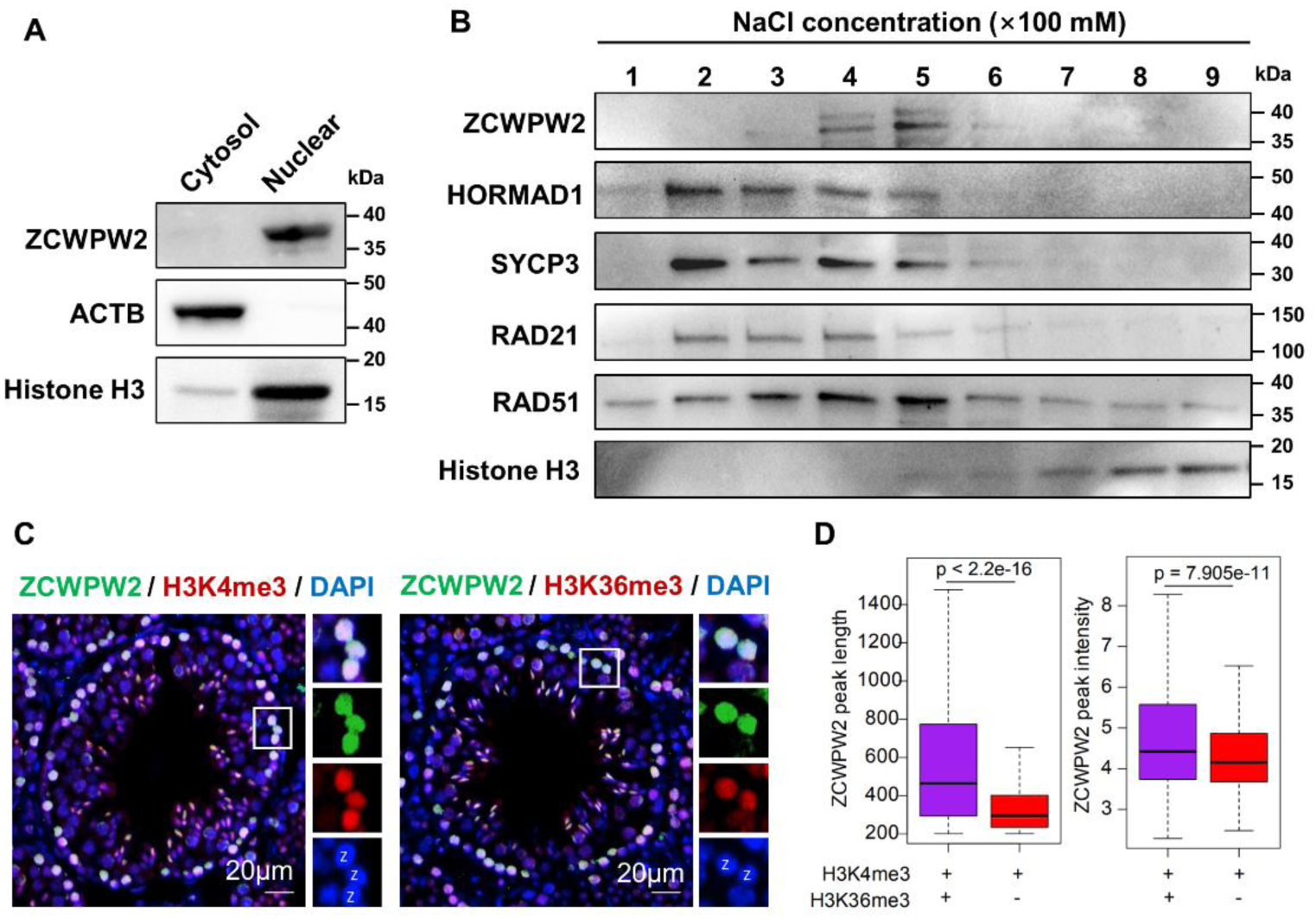
The expression and localization of ZCWPW2 in mice testes. **A.** Western bolting of ZCWPW2 in the cytoplasmic and nuclear fractions of PD12 wild type testes shows that ZCWPW2 localized in the nuclei. Histone H3 was used as the marker of nuclear fractions, and β-Actin was used as the marker of cytoplasmic fractions. **B.** Chromatin-binding assay of ZCWPW2 and indicated meiotic proteins. Chromatin fractions from wild type mouse testes were extracted and washed with increasing concentrations of NaCl (100–900 mM). Bound proteins were analyzed by Western blotting using antibodies against ZCWPW2, HORMAD1, SYCP3, RAD21, RAD51, and Histone H3. ZCWPW2 was detected in the high-salt eluates (400–500 mM NaCl), indicating its strong chromatin-binding activity. Histone H3 served as a control for chromatin-associated proteins. **C.** Immunostaining of adult mouse testicular sections showing co-localization of ZCWPW2 (green) with H3K4me3 (red, left panel) and H3K36me3 (red, right panel). Nuclei were counterstained with DAPI. Insets show magnified views of representative zygotene spermatocytes (labeled as Z). Scale bar, 20 μm. **D.** Boxplots comparing the difference of peak length and intensity between two ZCWPW2 groups shown in (Fig.1E). Wilcoxon rank sum test is used.

**Supplementary Figure S3.**
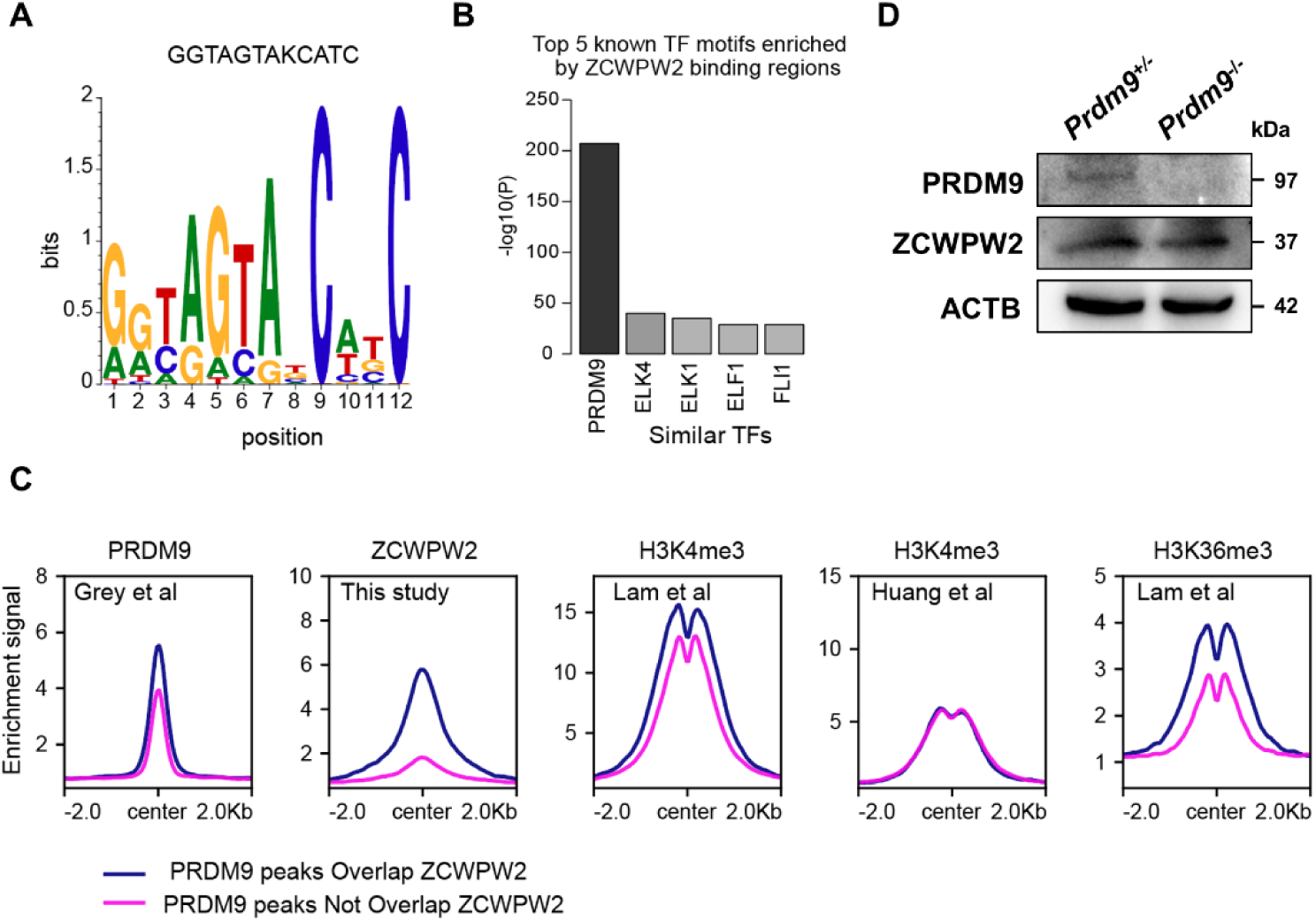
Genome-wide properties of ZCWPW2-associated binding sites. **A.** The rank-first de novo binding motif of ZCWPW2. **B.** Bar plot showing the similarity of ZCWPW2 binding motifs with those of other transcription factors. **C.** Profile plot showing the PRDM9, ZCWPW2, H3K4me3 and H3K36me3 signals on all the PRDM9 peaks in wild type testes. The PRDM9 peaks are classified into two groups including overlap with ZCWPW2 peaks (Group I) and without ZCWPW2 peaks (Group II). **D.** Western blot of PRDM9, ZCWPW2 in testes of wild type and *Prdm9^−/−^* mice at PD42. β-Actin was used as the loading control. PD, postnatal day.

**Supplementary Figure S4.**
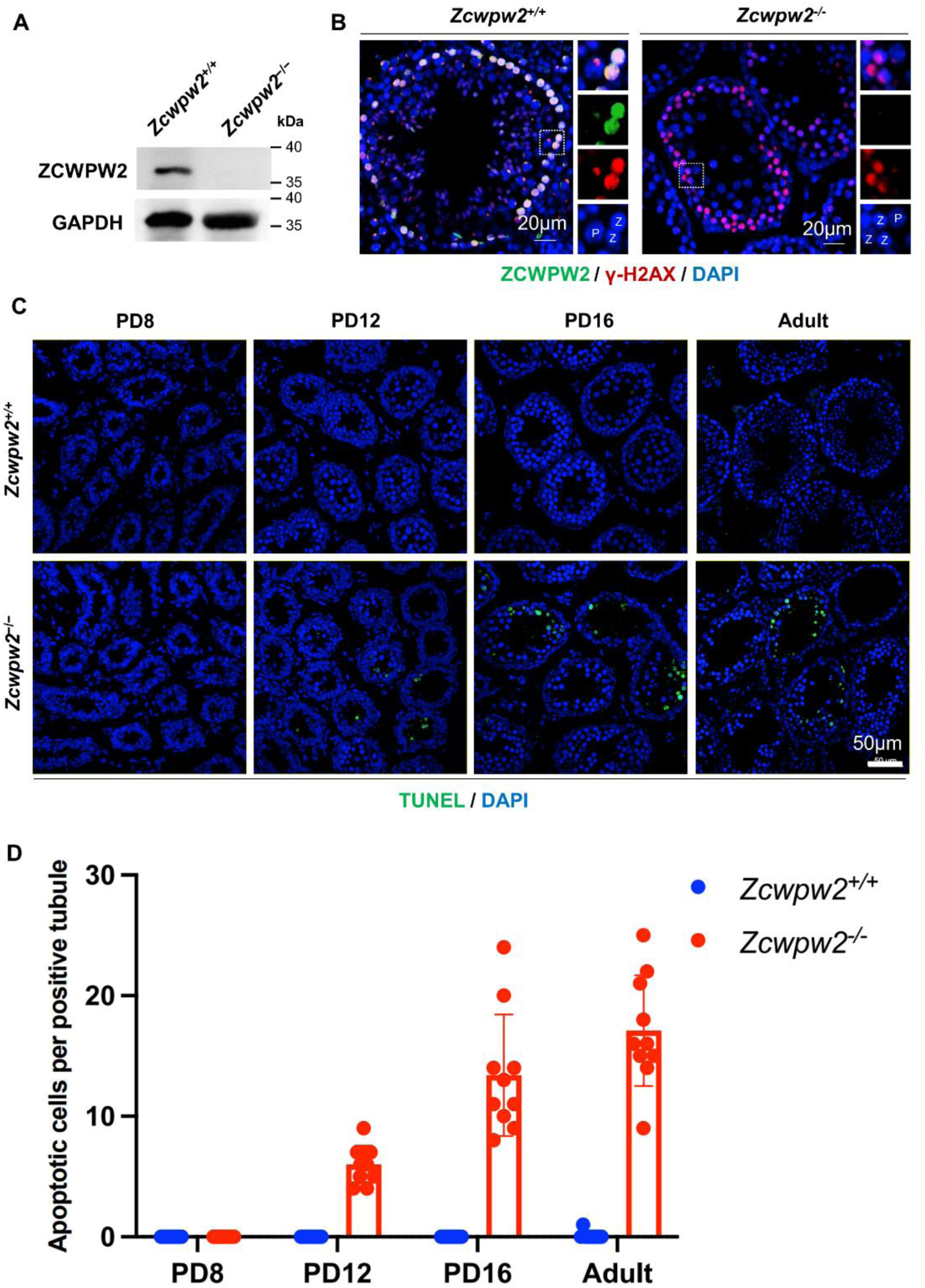
Deficiency of *Zcwpw2* results in apoptotic cells increased in *Zcwpw2^-/-^* mice testis. **A.** Western blotting analysis of ZCWPW2 in testes from PD14 *Zcwpw2^+/+^* and *Zcwpw2^-/-^* mice (n = 3). GAPDH served as a loading control. **B.** Double immunostaining for ZCWPW2 and γH2AX in adult *Zcwpw2^+/+^* and *Zcwpw2^-/-^* testes. DNA was counterstained with DAPI. Insets show magnified views of representative zygotene and pachytene spermatocytes (labeled as Z and P). Scale bar, 20 μm. **C.** TUNEL staining of seminiferous tubules showed that there were more apoptotic cells in *Zcwpw2^-/-^* mice compared with *Zcwpw2^+/+^* mice from PD12. The test was replicated three times using distinct biological samples. The scale bar is 50 μm. **D.** The number of apoptotic cells per seminiferous tubule in *Zcwpw2^+/+^* mice and *Zcwpw2^-/-^*mice. The data are shown as the mean ± SEM of three independent experiments using distinct biological samples. Each data point represents the number of apoptotic cells per seminiferous tubule.

**Supplementary Figure S5.**
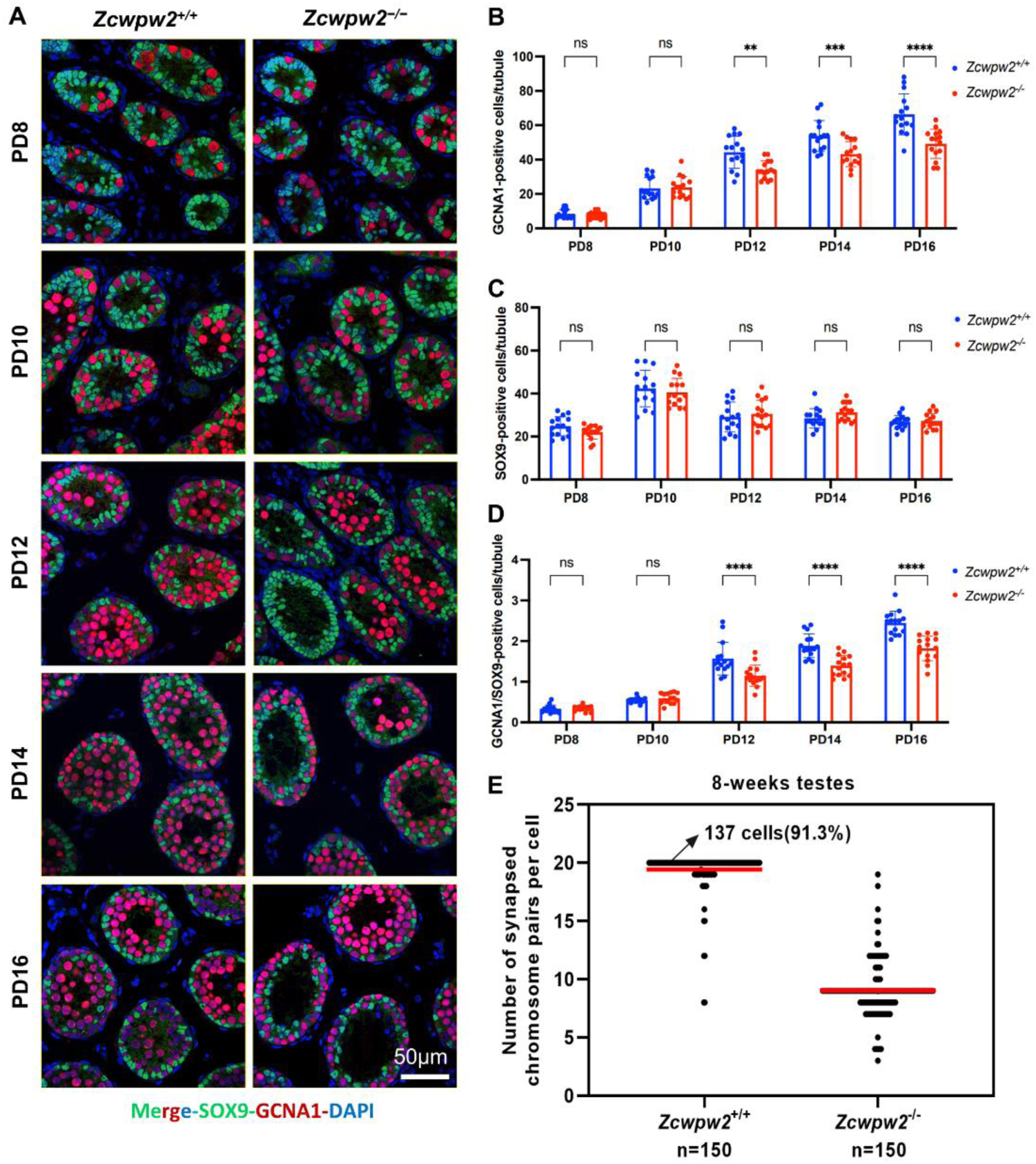
Deficiency of *Zcwpw2* results in germ cell loss in *Zcwpw2^-/-^* mice testis. **A.** Immunostaining for GCNA and SOX9 in *Zcwpw2^+/+^* and *Zcwpw2^-/-^* testes. DNA was counterstained with DAPI. Scale bar, 50 μm. **B.** The number of GCNA positive cells per seminiferous tubule in *Zcwpw2^+/+^* mice and *Zcwpw2^-/-^* mice. The data are shown as the mean ± SEM of three independent experiments using distinct biological samples. Each data point represents the number of GCNA positive cells per seminiferous tubule. **C.** The number of SOX9 positive cells per seminiferous tubule in *Zcwpw2^+/+^* mice and *Zcwpw2^-/-^* mice. The data are shown as the mean ± SEM of three independent experiments using distinct biological samples. Each data point represents the number of SOX9 positive cells per seminiferous tubule. **D.** The number of GCNA/SOX9 positive cells per seminiferous tubule in *Zcwpw2^+/+^* mice and *Zcwpw2^-/-^* mice. The data are shown as the mean ± SEM of three independent experiments using distinct biological samples. Each data point represents the number of GCNA/SOX9 positive cells per seminiferous tubule. **E.** The numbers of synapsed chromosome pairs in *Zcwpw2^+/+^* and *Zcwpw2^-/-^* spermatocytes. In *Zcwpw2^-/-^* spermatocytes, the average number of synapsed chromosome pairs was 8.

**Supplementary Figure S6.**
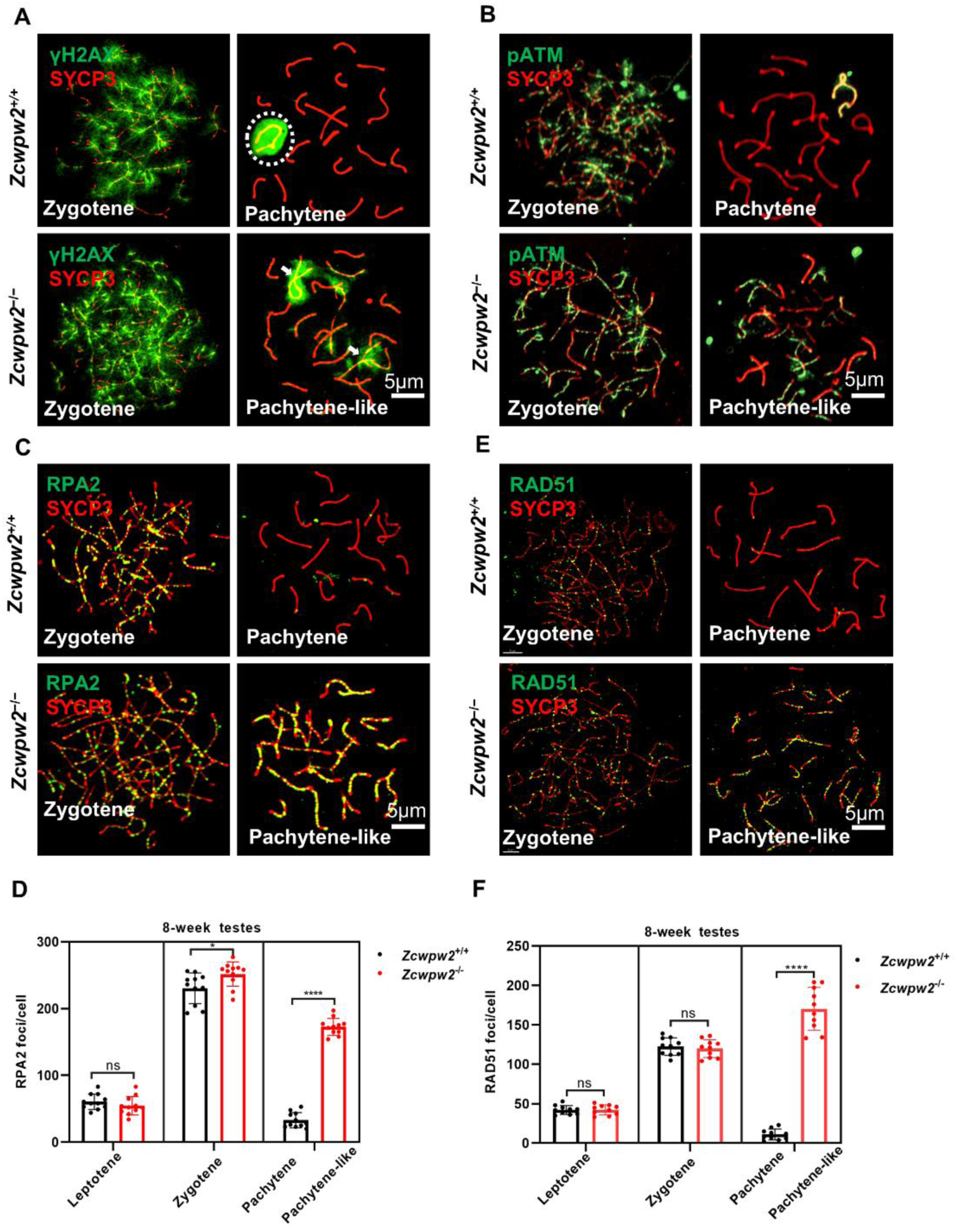
*Zcwpw2* knockout result in meiotic recombination defects in spermatocytes. **A.** Chromosome spreads of spermatocytes from the testes of adult *Zcwpw2^+/+^* and *Zcwpw2^-/-^*males were immunostained for the DSB marker γH2AX (green) and SYCP3 (red). Representative images are shown for spermatocytes at zygotene, pachytene (circle indicates the XY body), and pachytene-like (arrows indicate γH2AX signals) stages of the two genotypes. All experiments were performed on adult mice (6-8 weeks) with n = 3 for each genotype. Scale bar, 5 μm. **B.** Chromosome spreads of spermatocytes from the testes of adult *Zcwpw2^+/+^* and *Zcwpw2^-/-^* males were immunostained for the DSB marker pATM (green) and SYCP3 (red). Representative images are shown for spermatocytes at zygotene, pachytene, and pachytene-like stages of the two genotypes. Scale bar, 5 μm. **C.** Chromosome spreads of spermatocytes from the testes of adult *Zcwpw2^+/+^* and *Zcwpw2^-/-^*males were immunostained for RPA2 (green) and SYCP3 (red). Representative images of spermatocytes at zygotene and pachytene in *Zcwpw2^+/+^* and at zygotene and pachytene-like stages in *Zcwpw2^-/-^* are shown. Scale bar, 5 μm. **D.** Each dot represents the number of RPA2 foci per cell, with black dots indicating *Zcwpw2^+/+^* spermatocytes and red dots indicating *Zcwpw2^-/-^* spermatocytes. Solid lines show the mean and the SD of foci number in each group of spermatocytes. P values were calculated by Student’s t-test. N represents the number of cells. **E.** Chromosome spreads of spermatocytes from the testes of adult *Zcwpw2^+/+^* and *Zcwpw2^-/-^*males were immunostained for RAD51 (green) and SYCP3 (red). Representative images of spermatocytes at zygotene and pachytene in *Zcwpw2^+/+^* and at zygotene and pachytene-like stages in *Zcwpw2^-/-^* are shown. Scale bar, 5 μm. **F.** Each dot represents the number of RAD51 foci per cell, with black dots indicating *Zcwpw2^+/+^* spermatocytes and red dots indicating *Zcwpw2^-/-^* spermatocytes. Solid lines show the mean and SD of the number of foci in each group of spermatocytes. P values were calculated by Student’s t-test.

**Supplementary Figure S7.**
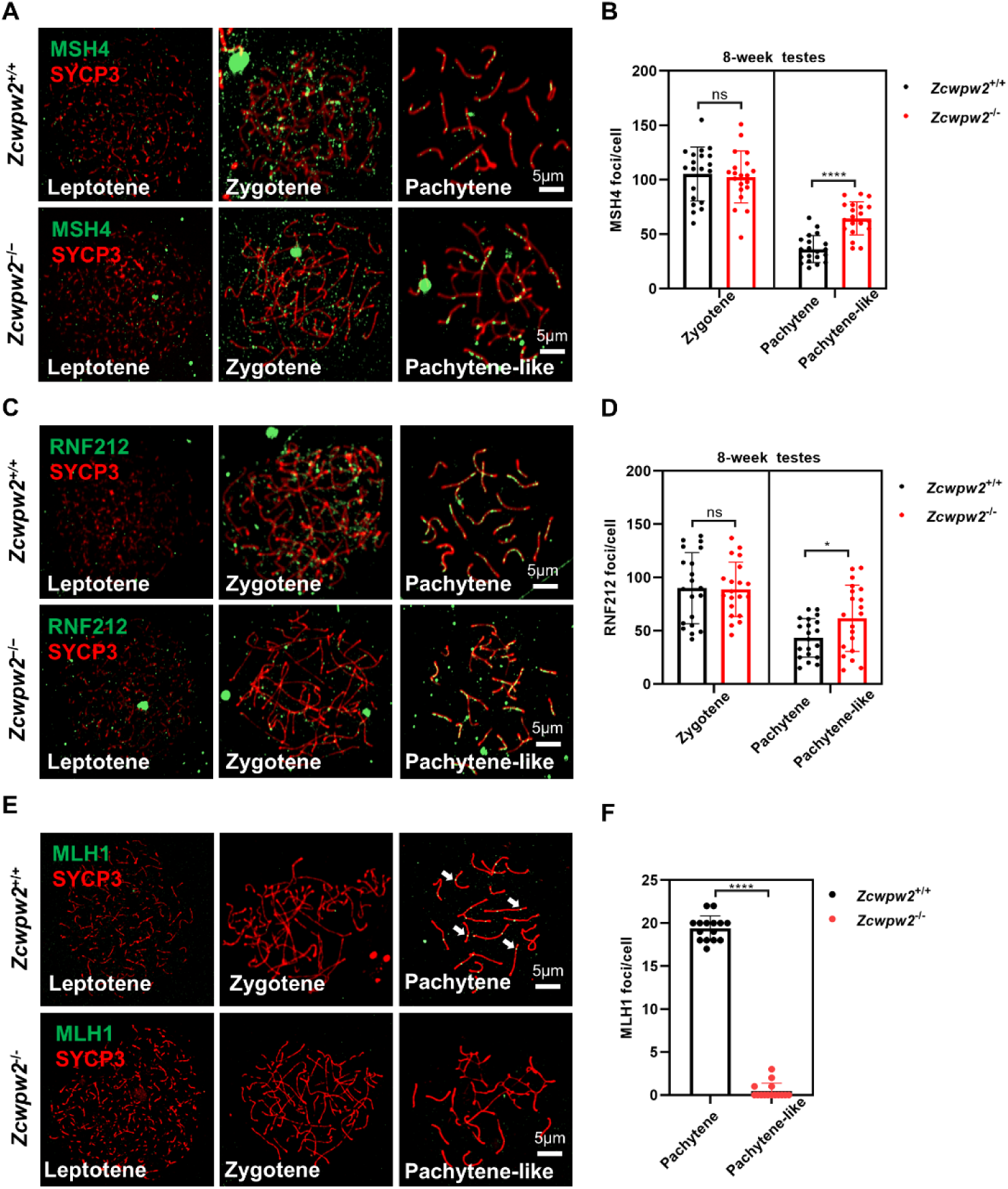
*Zcwpw2* deficiency impairs meiotic recombination in spermatocytes. **A.** *Zcwpw2^+/+^* and *Zcwpw2^-/-^* spermatocytes immunostained for MSH4 (green) and SYCP3 (red). Representative images are shown for spermatocytes at leptotene, zygotene, pachytene and pachytene-like stages of the two genotypes. Scale bar, 5 μm. **B.** Each dot represents the number of MSH4 foci per cell, with black dots indicating *Zcwpw2^+/+^* spermatocytes and red dots indicating *Zcwpw2^-/-^* spermatocytes. Solid lines show the mean and SD of the foci number for each group of spermatocytes. **C.** *Zcwpw2^+/+^* and *Zcwpw2^-/-^* spermatocytes immunostained for RNF212 (green) and SYCP3 (red). Representative images are shown for spermatocytes at leptotene, zygotene, pachytene and pachytene-like stages of the two genotypes. Scale bar, 5 μm. **D.** Each dot represents the number of RNF212 foci per cell, with black dots indicating *Zcwpw2^+/+^* spermatocytes and red dots indicating *Zcwpw2^-/-^* spermatocytes. Solid lines show the mean and SD of the number of foci for each group of spermatocytes. **E.** *Zcwpw2^+/+^* and *Zcwpw2^-/-^* spermatocytes were immunostained for MLH1 (green) and SYCP3 (red). *Zcwpw2^-/-^* spermatocytes lack MLH1 signal. Representative images are shown for spermatocytes at leptotene, zygotene, pachytene and pachytene-like stages of the two genotypes. Scale bar, 5 μm. **F.** Each dot represents the number of MLH1 foci per cell, with black dots indicating *Zcwpw2^+/+^* spermatocytes and red dots indicating *Zcwpw2^-/-^* spermatocytes. Solid lines show the mean and SD of the number of foci for each group of spermatocytes.

**Supplementary Figure S8.**
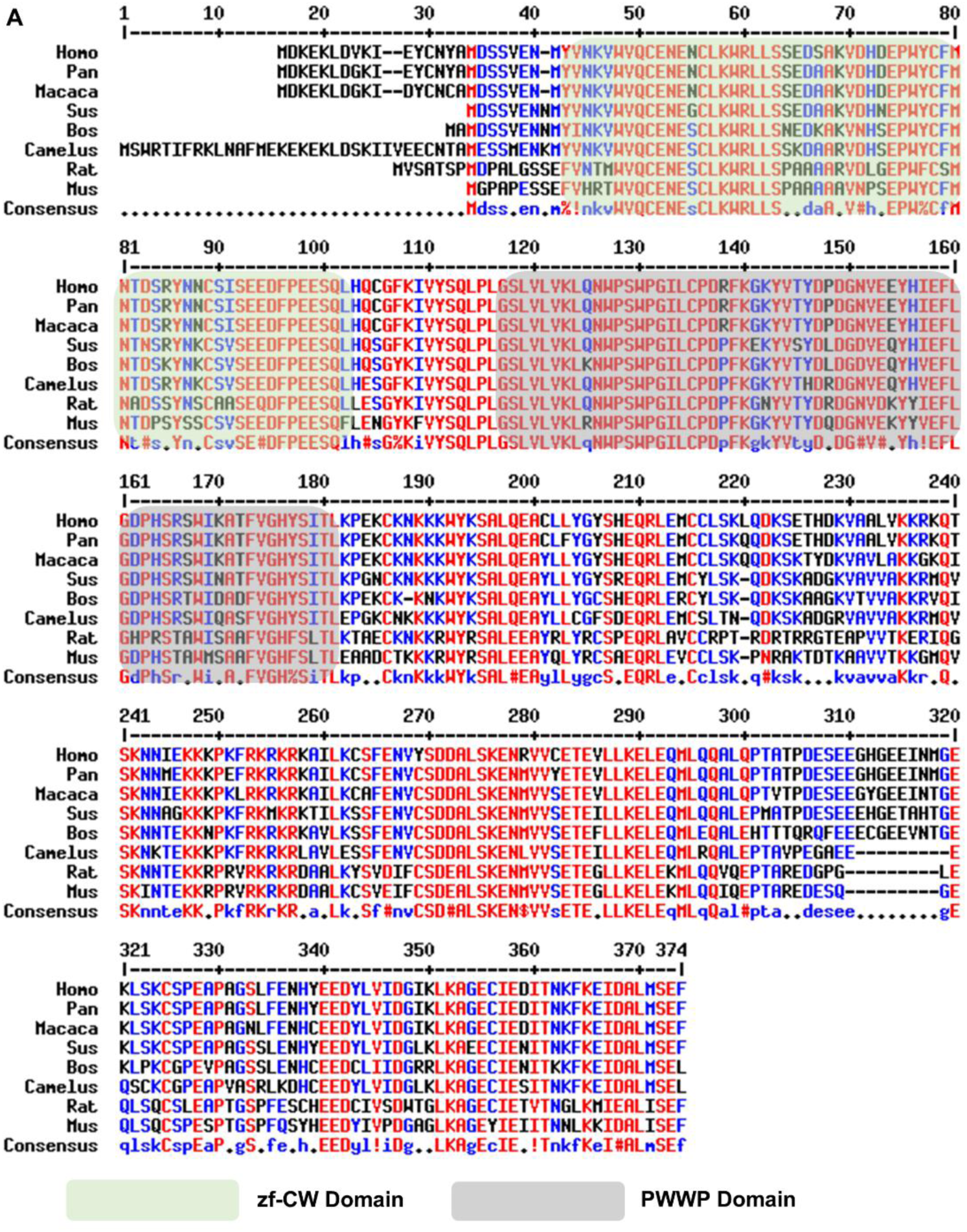
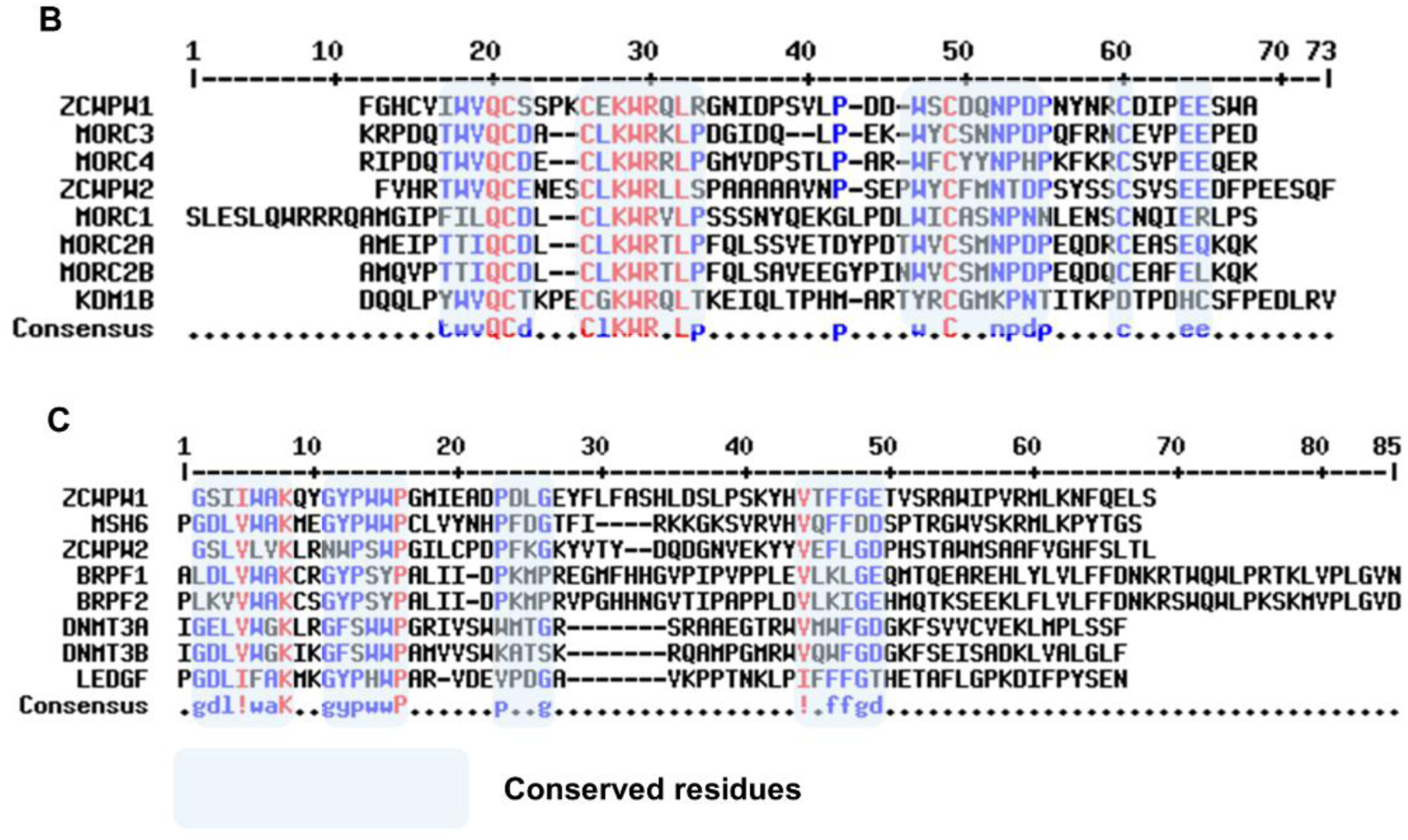
Sequence alignment of ZCWPW2 between different species. **A.** Sequence alignment of ZCWPW2 between different species, such as humans, non-human primates, rodents, and ruminants. Conserved regions of ZCWPW2 are indicated by green shaded areas for the zf-CW domain and grey shaded areas for the PWWP domain. **B.** Sequence alignment of zf-CW domain between different proteins. Conserved amino acids of zf-CW domain are indicated by green shaded areas. **C.** Sequence alignment of PWWP domain between different proteins. Conserved amino acids of PWWP domain are indicated by green shaded areas.

**Supplementary Figure S9.**
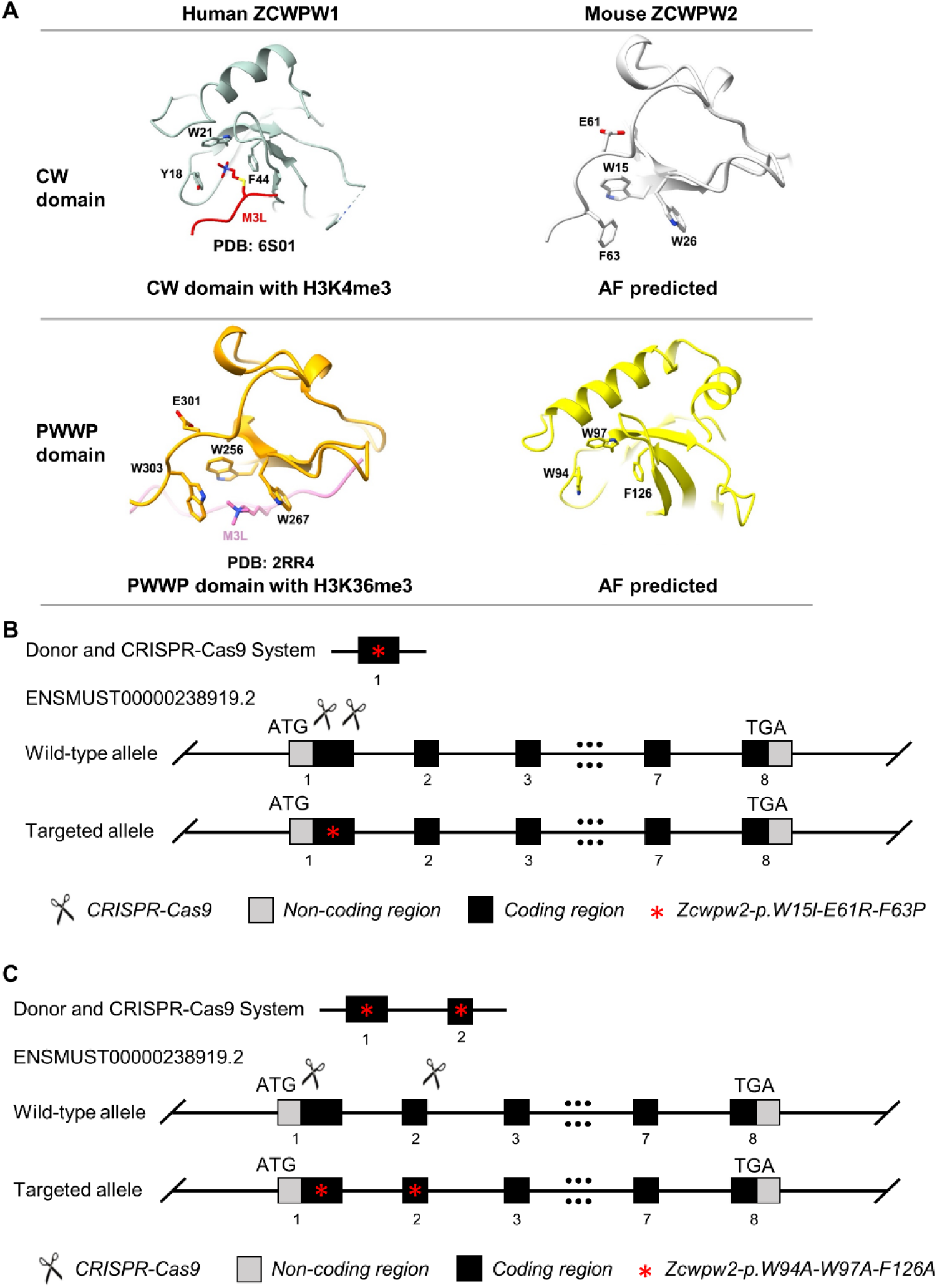
Generation of *Zcwpw2* reader-dead-mutant mice. **A.** Structure comparison of the human ZCWPW1 zf-CW domain and PWWP domain with mouse ZCWPW2 zf-CW domain and PWWP domain. Left panel was superpositions of the free zf-CW domain and PWWP domain, showing the residues forming the aromatic cage and the H3K4me3 or H3K36me3 peptide. Right panel was the AlphaFold3 predicting a homologous structure in the mouse ZCWPW2 zf-CW domain and PWWP domain. **B.** Schematic representation of the CRISPR/Cas9 genome editing system for generating *Zcwpw2^CW-KI^* mice. **C.** Schematic representation of the CRISPR/Cas9 genome editing system for generating *Zcwpw2^PWWP-KI^* mice.

**Supplementary Figure S10.**
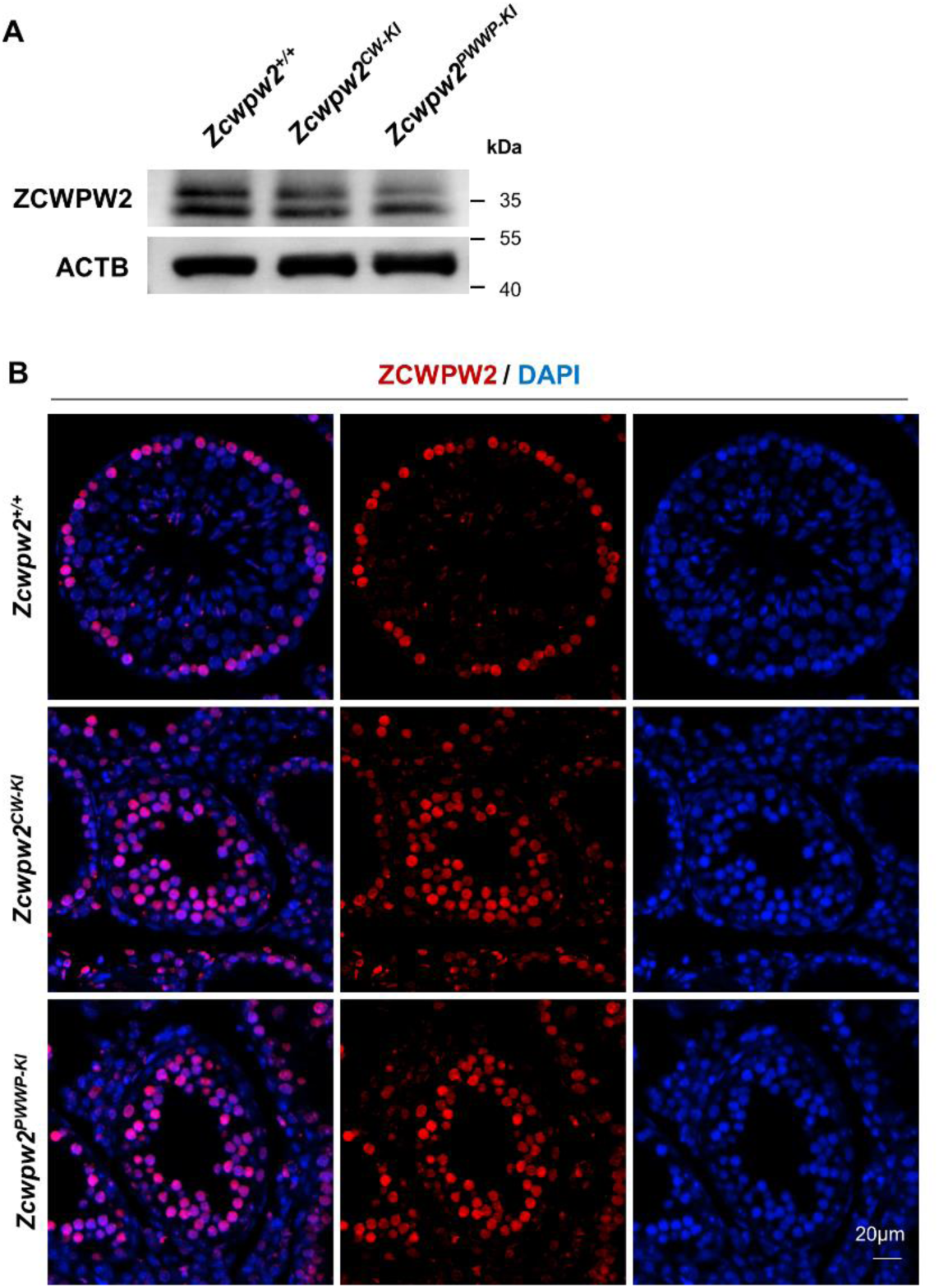
The expression and localization of ZCWPW2 were normal in *Zcwpw2* reader dead mutant mice. **A.** Western blotting analysis of ZCWPW2 in testes from PD14 *Zcwpw2^+/+^*, *Zcwpw2^CW-KI^* and *Zcwpw2^PWWP-KI^* mice. ACTB served as a loading control. **B.** Immunostaining for ZCWPW2 in adult *Zcwpw2^+/+^*, *Zcwpw2^CW-KI^* and *Zcwpw2^PWWP-KI^* mouse testes. DNA was counterstained with DAPI. Scale bar, 20 μm.

**Supplementary Figure S11.**
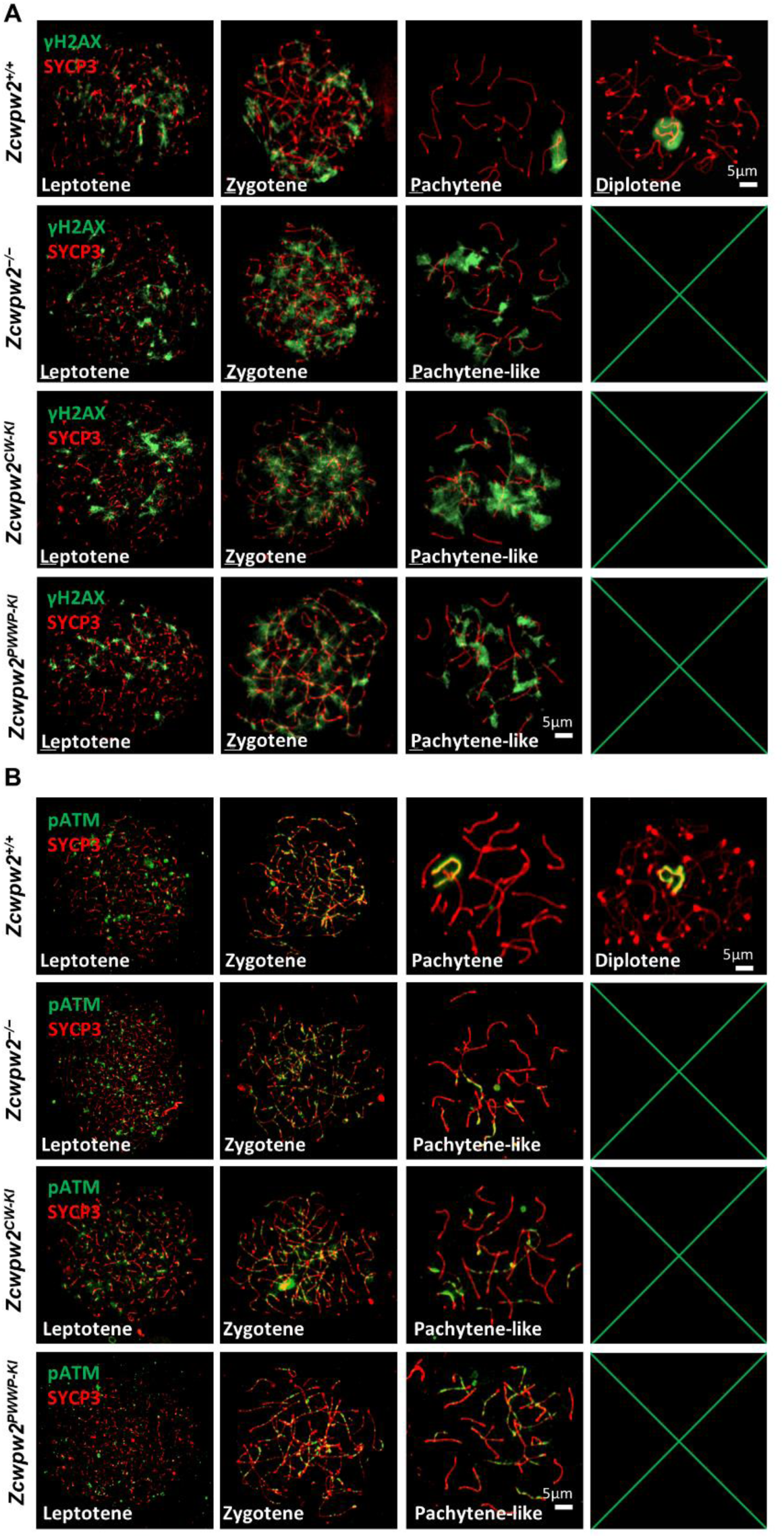
Meiotic DSB repair defects in *Zcwpw2* reader-dead-mutant mice. **A.** Chromosome spreads of spermatocytes from the testes of adult *Zcwpw2^+/+^*, *Zcwpw2^−/−^*, *Zcwpw2^CW-KI^*, and *Zcwpw2^PWWP-KI^* males were immunostained for the DSB marker proteins γH2AX (green) and SYCP3 (red). Representative images are shown for spermatocytes at leptotene, zygotene, pachytene, pachytene-like, and diplotene stages across the four genotypes. Scale bar, 5 μm. **B.** Chromosome spreads of spermatocytes from the testes of adult *Zcwpw2^+/+^*, *Zcwpw2^−/−^*, *Zcwpw2^CW-KI^*, and *Zcwpw2^PWWP-KI^* males were immunostained for pATM (green) and SYCP3 (red). Representative images are shown for spermatocytes at leptotene, zygotene, pachytene, pachytene-like, and diplotene stages across the four genotypes. Scale bar, 5 μm.

**Supplementary Figure S12.**
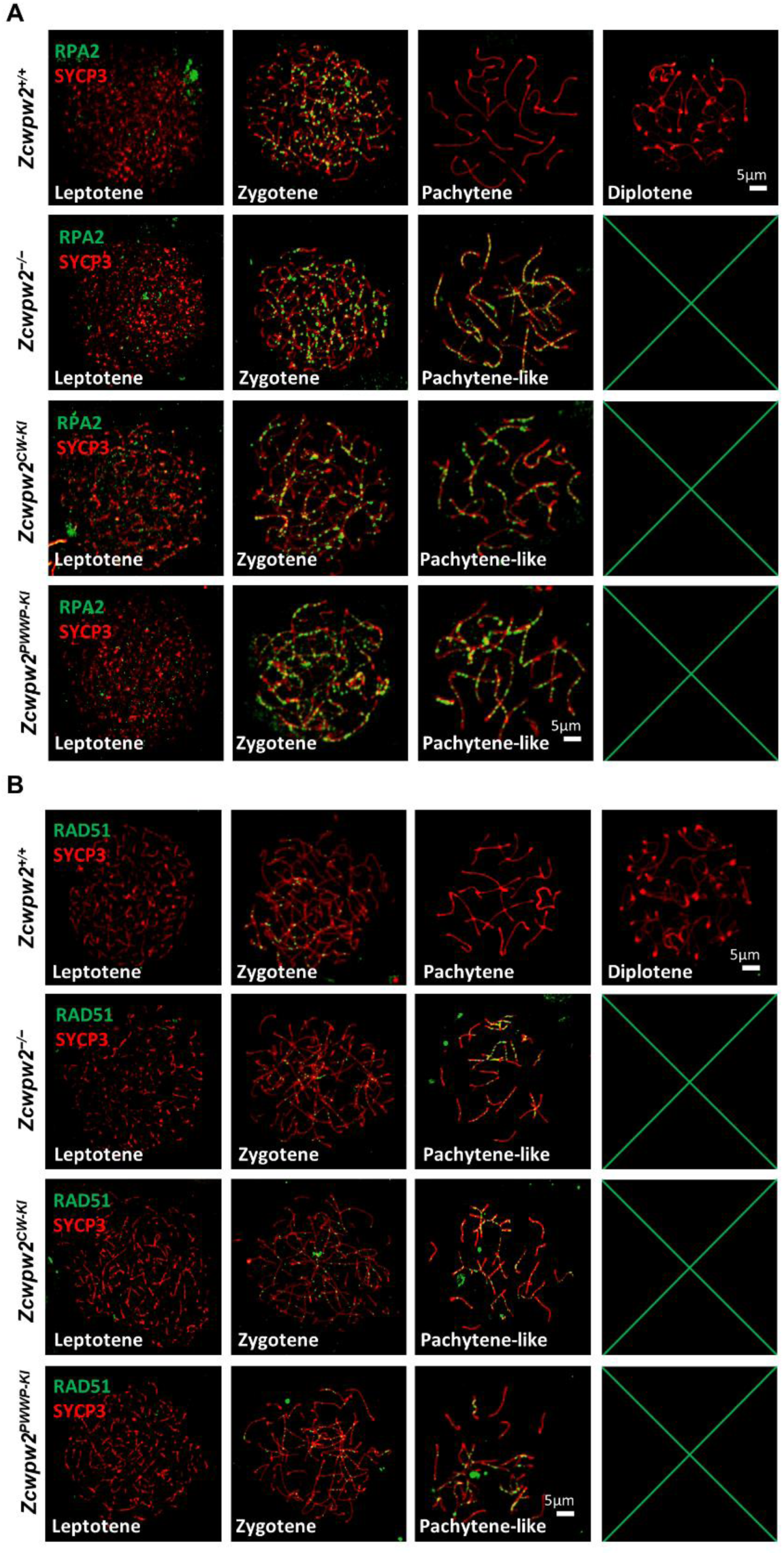
Meiotic recombination defects (RPA2 and RAD51) in *Zcwpw2* reader-dead-mutant mice. **A.** Chromosome spreads of spermatocytes from the testes of adult *Zcwpw2^+/+^*, *Zcwpw2^−/−^*, *Zcwpw2^CW-KI^*, and *Zcwpw2^PWWP-KI^* males were immunostained for RPA2 (green) and SYCP3 (red). Representative images are shown for spermatocytes at leptotene, zygotene, pachytene, pachytene-like, and diplotene stages across the four genotypes. Scale bar, 5 μm. **B.** Chromosome spreads of spermatocytes from the testes of adult *Zcwpw2^+/+^*, *Zcwpw2^−/−^*, *Zcwpw2^CW-KI^*, and *Zcwpw2^PWWP-KI^* males were immunostained for RAD51 (green) and SYCP3 (red). Representative images are shown for spermatocytes at leptotene, zygotene, pachytene, pachytene-like, and diplotene stages across the four genotypes. Scale bar, 5 μm.

**Supplementary Figure S13.**
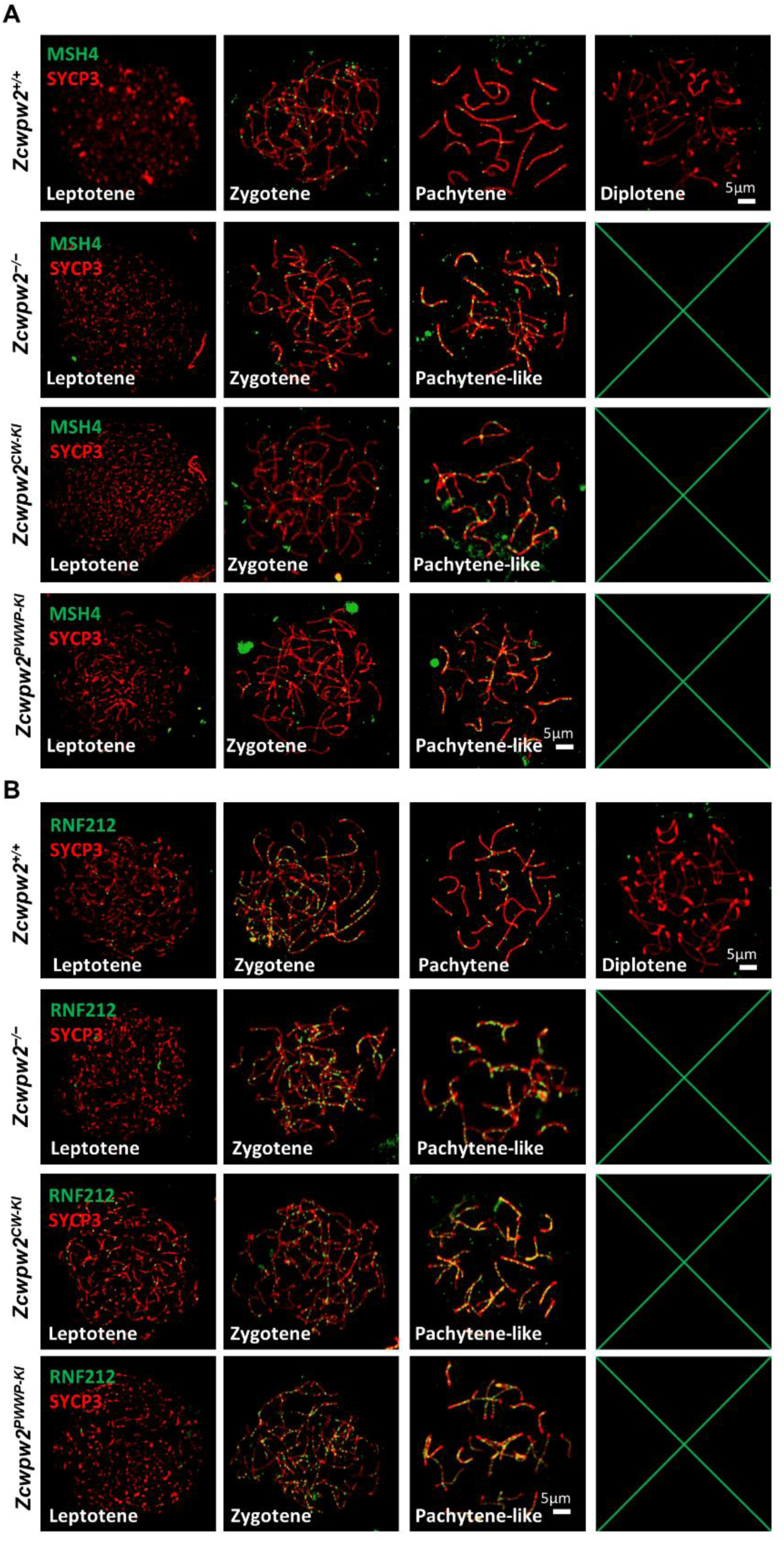
Meiotic recombination defects (MSH4 and RNF212) in *Zcwpw2* reader-dead-mutant mice. **A.** Chromosome spreads of spermatocytes from the testes of adult *Zcwpw2^+/+^*, *Zcwpw2^−/−^*, *Zcwpw2^CW-KI^*, and *Zcwpw2^PWWP-KI^* males were immunostained for MSH4 (green) and SYCP3 (red). Representative images are shown for spermatocytes at leptotene, zygotene, pachytene, pachytene-like, and diplotene stages across the four genotypes. Scale bar, 5 μm. **B.** Chromosome spreads of spermatocytes from the testes of adult *Zcwpw2^+/+^*, *Zcwpw2^−/−^*, *Zcwpw2^CW-KI^*, and *Zcwpw2^PWWP-KI^* males were immunostained for RNF212 (green) and SYCP3 (red). Representative images are shown for spermatocytes at leptotene, zygotene, pachytene, pachytene-like, and diplotene stages across the four genotypes. Scale bar, 5 μm.

**Supplementary Figure S14.**
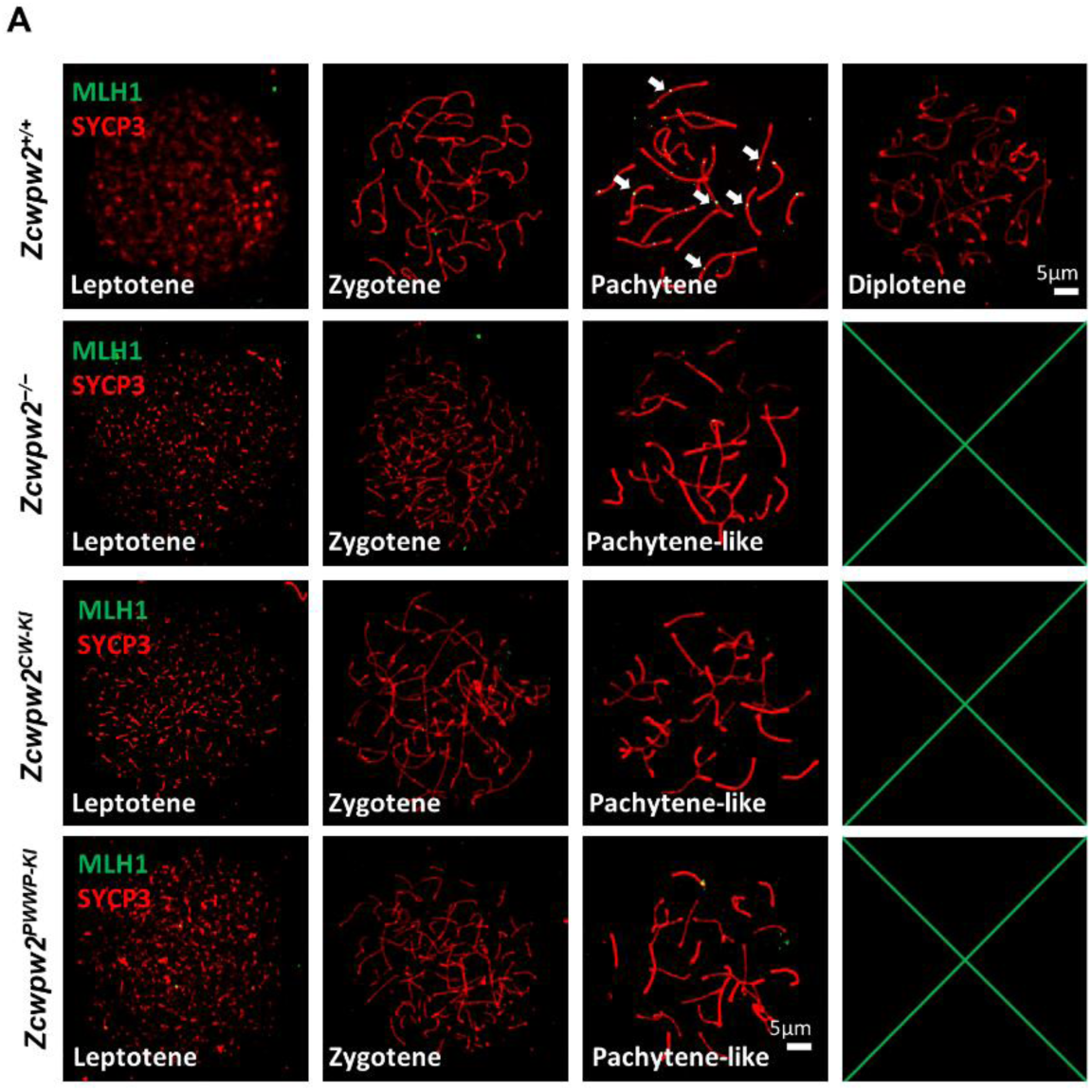
Meiotic recombination defects (MLH1) in *Zcwpw2* reader-dead-mutant mice. **A.** Chromosome spreads of spermatocytes from the testes of adult *Zcwpw2^+/+^*, *Zcwpw2^−/−^*, *Zcwpw2^CW-KI^*, and *Zcwpw2^PWWP-KI^* males were immunostained for MLH1 (green) and SYCP3 (red). Representative images are shown for spermatocytes at leptotene, zygotene, pachytene (arrows indicate the MLH1 signal), pachytene-like, and diplotene stages across the four genotypes. Scale bar, 5 μm.

**Supplementary Figure S15.**
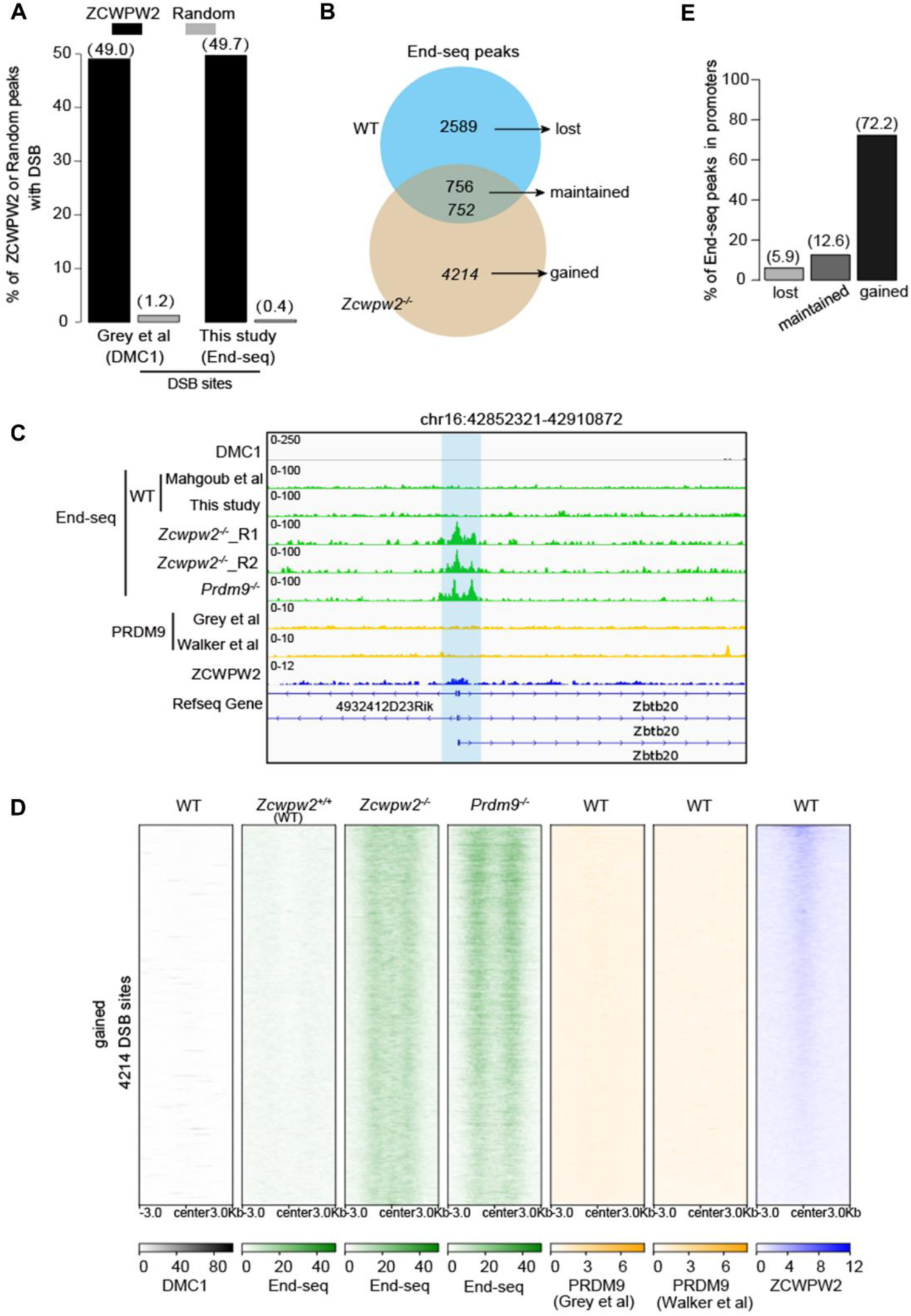
Genome-wide properties of DSB in *Zcwpw2^-/-^* spermatocytes. **A.** Bar plot showing the percentage of ZCWPW2 peaks or the random regions overlapping DMC1 or End-seq peaks. **B.** Venn plot illustrating the alteration in the count of End-seq peaks between in wild type and *Zcwpw2^-/-^* testis. The End-seq peaks are classified into three groups including lost, maintained and gained peaks. **C.** Genome browser view of DMC1, PRDM9 and ZCWPW2 signal in wild type testes, Enq-seq signal in wild type, *Zcwpw2^-/-^* and *Prdm9^-/-^* testis. The blue shadows indicate the ZCWPW2 and PRDM9 binding regions with DMC1 and End-seq peaks in *Zcwpw2^-/-^* and *Prdm9^-/-^* testes, but lost in wild type testes. **D.** Heatmap showing the difference of DSB signal on the gained END-seq peaks in *Zcwpw2^-/-^* and *Prdm9^-/-^* testis. DMC1, PRDM9, and ZCWPW2 signals in wild-type testes are also shown. **E.** Bar plot comparing the percentage of three groups of End-seq peaks in promoter regions.

**Supplementary Figure S16.**
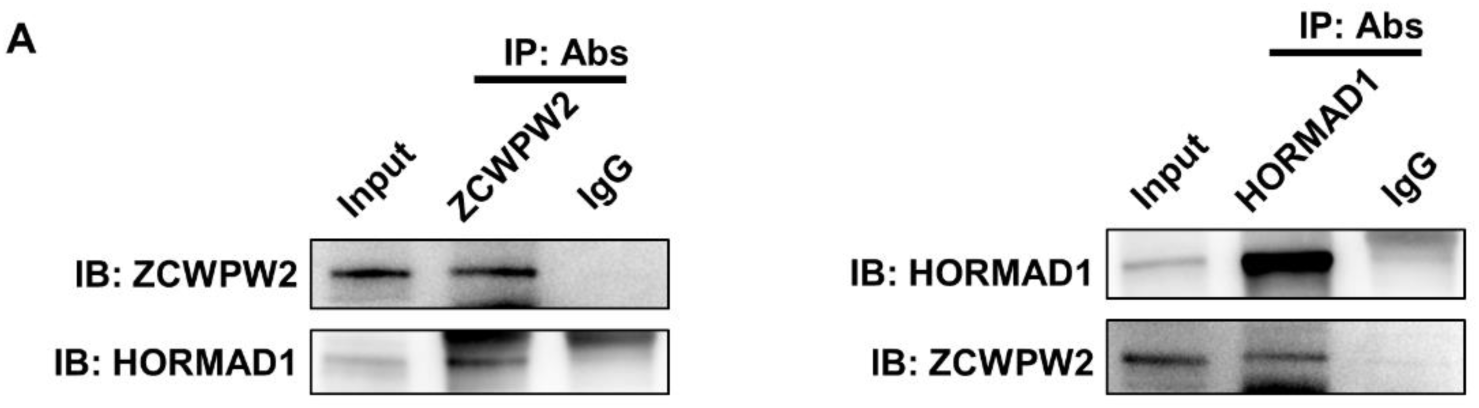
ZCWPW2 physically interacts with HORMAD1 in mouse testes. **A.** Co-IP analysis of the interaction between ZCWPW2 and HORMAD1 in wild type mouse testes. Data are representative of three independent experiments.

## Supplementary Tables

**Supplementary Table1:** ZCWPW2_peaks_mouse_mm10.

**Supplementary Table2:** ZCWPW2_Data_resource.

**Supplementary Table3:** Antibodies are used in this study.

**Supplementary Table4:** Quantification of fluorescence images in this study.

## Methods

### Mice

The *Prdm9* knockout mice (*Prdm9^-/-^*) were generated in our previous study[49, 50]. The mouse *Zcwpw2* gene (GenBank accession number: NM_001378466.1; Ensembl: ENSMUST00000238919) is located on mouse chromosome 9 and comprises 8 exons, with the ATG start codon in exon 1 and the TAG stop codon in exon 8. The *Zcwpw2* knockout mice (*Zcwpw2^-/-^*) were generated by deleting exon 2 to exon 6 genomic DNA fragment using the CRISPR/Cas9-mediated genome editing system in a C57BL/6 genetic background (Cyagen Biosciences, Suzhou, China). For the *Zcwpw2* knockin model, the "3xFLAG tag-HA tag-2xGGGGS-APEX2" cassette was inserted upstream of *Zcwpw2* gene TAG stop codon in a C57BL/6J genetic background (Cyagen Biosciences, Suzhou, China).

The *Zcwpw2* knock-in H3K4me3-reader-dead mutant mice were generated by mutating three sites. The W15I (TGG to ATC), E61R (GAG to AGG) and F63P (TTC to CCT) point mutations were introduced into exon 1 (Shanghai Model Organisms Center, Inc). The *Zcwpw2* knock-in H3K36me3-reader-dead mutant mice were generated by mutating three sites. The W94A (TGG to GCC) point mutation was introduced into exon 1. The W97A (TGG to GCC) and F126A (TTT to GCC) point mutations were introduced into exon 2 (GemPharmatech, Nanjing, China). The *Zcwpw2* mutant founders were genotyped by PCR followed by DNA sequencing analysis. The homozygous mutant mice were generated by intercross of heterozygous mutants. Genotyping was performed by PCR amplification of genomic DNA extracted from mouse tails. PCR primers for the *Zcwpw2* knockout allele were Forward: 5’- TGC CTA GCA TAC TTT GCT TCA TCT-3’ and Reverse: 5’- ATG ACC CAC TAA CCT TTT ACT CCT-3’, yielding a 628 bp fragment. PCR primers for the *Zcwpw2* wild-type allele were Forward: 5’- TCA TGG TTT TAG TTTCTTTGCCTGG -3’ and Reverse: 5’- ATG ACC CAC TAA CCT TTT ACT CCT -3’, yielding a 521 bp fragment.

PCR primers for the *Zcwpw2* knockin allele were Forward: 5’- GCT TTG GAG GAC CTT CTG TAT T -3’ and Reverse: 5’- GCA CTC ACA GTT GGG TAA GAC TTT C -3’, yielding a 359 bp fragment. PCR primers for the *Zcwpw2* wild-type allele were Forward: 5’- GCT TTG GAG GAC CTT CTG TAT T-3’ and Reverse: 5’- GCC TAA ATA CTC ATG AAC AGC ACC -3’, yielding a 514 bp fragment.

PCR primers for the *Zcwpw2* CW mutant were Forward: 5’- CTT CTC TGC CAG GAG CGT AGG A -3’ and Reverse: 5’- ACA CAC ACA AAC ACA TGT CCA T -3’, yielding a 620 bp fragment. Sequencing primer for the *Zcwpw2* CW mutant was Forward: 5’- TTC TGA GAA ACA AGA CAA AA -3’.

PCR primers for the *Zcwpw2* PWWP mutant were Forward: 5’- CAG AGC ACT TGT TAT GGT CAT TTA CGT G -3’ and Reverse: 5’- CCT AGC CAG TGT GAG TGA GGC TAT T -3’, yielding a 767 bp fragment. Sequencing primer for the *Zcwpw2* PWWP mutant was Forward: 5’- CAG AGC ACT TGT TAT GGT CAT TTA CGT G -3’.

All mice were maintained with free access to water and food at the specific pathogen-free (SPF) facility under a 12-h light/dark cycle. All experimental protocols were approved by the regional ethics committee of Shandong University.

### Production of antibody

Antibodies to mouse ZCWPW2 were produced by MabStar antibody technology (Wuhan) Co., Ltd. Antibodies to mouse PRDM9 were produced by Kwinbon Technology Co., Ltd. (Beijing, China). Briefly, a complementary DNA (cDNA) fragment encoding amino acids 414 to 537 of mouse *Prdm9* and amino acids 151 to 331 of mouse *Zcwpw2* were chosen because of their hydrophilicity and immunogenicity. Then they were inserted into the p-ET-32a + vector (EMD Millipore) and transfected into BL21-CodonPlus (DE3) Escherichia coli cells. The cells were cultured at 37°C overnight and induced by addition of 0.2 mM isopropyl-1-thio-β -D-galactoside (Sigma-Aldrich) for 4 hours at 28°C. Cells were harvested by centrifugation and disrupted by sonication, and the soluble homogenates were purified by Ni-nitrilotriacetic acid (NI-NTA) Agarose (Qiagen) according to the manufacturer’s instructions. The proteins were dialyzed in phosphate-buffered saline (PBS) and used to immunize rabbits, and the antiserums were affinity-purified on antigen-coupled CNBr-activated agarose (GE Healthcare).

### Fertility test

Sexually mature *Zcwpw2^+/+^* and *Zcwpw2^-/-^*mice (8 weeks old) were housed individually with three wildtype females each. Mating success was verified by checking for vaginal plugs every morning. Once a plug was confirmed, the female was separated and housed alone, and the number of offspring per litter was recorded to evaluate fertility. A minimum of three *Zcwpw2^+/+^* and *Zcwpw2^-/-^*males were used in the fecundity assessments.

### Tissue collection and histological analysis

Following euthanasia, testes and epididymides from at least three mice per genotype were promptly collected and fixed overnight in Bouin’s Fixative Solution (Sigma-Aldrich, HT10132) at 4°C. After fixation, the specimens were dehydrated using a graded ethanol series, cleared with xylene, and embedded in paraffin. The resulting blocks were sectioned at a thickness of 5 μm and mounted onto glass slides. For histological examination, the sections were deparaffinized in xylene, rehydrated through a descending ethanol series to distilled water, and stained with hematoxylin (Beyotime, C0105M). Images were captured using a fluorescence microscope (BX53, Olympus).

### Chromosome spreading

After euthanasia, mouse testes were dissected, and the tunica albuginea was removed. The remaining tissue was incubated in a hypotonic solution (30 mM Tris, 50 mM sucrose, 17 mM sodium citrate, 5 mM EDTA, 0.5 mM dithiothreitol [DTT], pH 9.2) for 30 minutes at room temperature. Subsequently, the seminiferous tubules were transferred to 100 mM sucrose, thoroughly dissociated, and spread onto slides pre-treated with 1% PFA. The slides were placed in a humid chamber, then air-dried at room temperature. Following drying, the sample areas were encircled with a histochemical pen, and the slides were stored at –80°C until further use.

### Immunofluorescence

Testes from at least three mice per genotype were immediately dissected and fixed in 4% paraformaldehyde (Solarbio, P1110) overnight at 4℃ after euthanasia. Following dehydration, embedding, sectioning, deparaffinization and rehydration, antigen retrieval was performed by boiling slides in Citrate Antigen Retrieval Solution (Beyotime, P0081) for 15 min, and cooling to RT. Sections were permeabilized with PBS (Gibco, C20012500BT) containing 0.1% Triton X-100 (Sigma, T8787) for 15 min and washed three times with PBS. Sections were blocked with 5% BSA (Amresco, NB200) for 1h at RT, then incubated with primary antibodies overnight at 4℃. After three PBS washes, secondary antibodies were added and incubated for 1h at RT. Following an additional three PBS washes, sections were mounted with DAPI-containing aqueous fluoroshield mounting medium (Abcam, ab104139). Antibodies used in this study were summarized in Supplementary Table S3.

### Microscopy

Immunostained slides were imaged by confocal microscopy (Andor Dragonfly spinning disc confocal microscope driven by Fusion Software). Projection images were then prepared using Photoshop (Adobe) software packages or Bitplane Imaris (version 8.1) software.

### Plasmid construction

cDNA for mouse *Zcwpw2, Hormad1, Iho1, Mei4*, and *Rec114* were synthesized by Wuhan GeneCreate Biological Engineering. The *Zcwpw2* and *Hormad1* were cloned into pCAG-GS vector for transient expression in cell lines. The *Iho1, Mei4,* and *Rec114* were cloned into pCAG-GS vector with a C-terminal fusion Flag-His tag for transient expression in cell lines. The integrity of all expression plasmids was verified by Sanger sequencing prior to use.

### Cell culture and transfection

HEK293T cells purchased from National Collection of Authenticated Cell Cultures were maintained in Dulbecco’s modified Eagle’s medium (DMEM) supplemented with 10% (v/v) fetal bovine serum and 1% penicillin-streptomycin at 37 °C in a 5% CO2 incubator. For transient transfection, cells were seeded into appropriate culture vessels and allowed to reach 70-80% confluence. Transfection was performed using X-tremeGENE HP DNA Transfection Reagent (Roche, #6366236001) according to the supplier’s protocol, with the medium being replaced 4-6 hours later. Cells were typically harvested 48 hours post-transfection for downstream analyses.

### Co-immunoprecipitation and western blotting

Total protein extracts were prepared from either mouse testes or transfected HEK293T cells. Tissue samples were minced and homogenized in ice-cold Pierce IP lysis buffer (25 mM Tris-HCl pH 7.4, 150 mM NaCl, 1 mM EDTA, 1% NP-40, and 5% glycerol; Thermo Scientific #87787) with protease inhibitors (Roche, #04693132001). For cell pellets, lysis was achieved by resuspending cells directly in the same buffer and pipetting gently. After incubation on ice for 30 min with intermittent vortexing, lysates were clarified by centrifugation at 12,000×g for 15 min at 4 °C. The supernatant was transferred into a new tube and incubated with primary antibody or control IgG with rotation overnight at 4 °C. Then, the antibodies were isolated by adsorption to Pierce protein A/G beads (Thermo Scientific, #88802) for 2 h. After washing, SDS loading buffer was added to the beads and boiled.

Samples, along with input controls, were resolved by SDS-PAGE using 10% or 4-20% gradient gels and electroblotted onto polyvinylidene difluoride (PVDF) membranes. Membranes were blocked for 1 h at room temperature with 5% (w/v) non-fat milk in TBST (Tris-buffered saline containing 0.1% Tween 20) and incubated with primary antibodies diluted in blocking buffer at 4 °C overnight. After extensive washing with TBST, membranes were probed with horseradish peroxidase-conjugated secondary antibodies for 1 h at room temperature. Immunoreactive signals were developed using enhanced chemiluminescence substrate and captured with a Bio-Rad ChemiDoc MP Imaging System. Band intensities were quantified with Image Lab software (Bio-Rad). Antibodies used in this study were summarized in Supplementary Table S3.

### Cytoplasmic and nuclear extraction

For native cytosolic and nuclear protein isolation from tissue, testes from wild-type mice were processed with a MinuteTM kit for frozen/fresh tissues (Invent Biotechnologies, NT-032). Briefly, 20-30 mg of tissue was immersed in 250 µL buffer A and chilled on ice, then homogenized with the supplied pestle. After centrifugation, the supernatant was retained as the cytosolic fraction. The remaining pellet was re-homogenized, suspended in additional buffer A, and left on ice to allow debris to settle. The supernatant was collected and centrifuged at low speed to pellet the nuclei. The pellet contains isolated nuclei. Antibodies used in this study were summarized in Supplementary Table S3.

### Sequential Salt Extractions

A series of 1× mRIPA buffers (100 mM Tris pH 8.0, 2% NP-40, 0.5% sodium deoxycholate) were prepared with NaCl concentrations ranging from 0 to 1000 mM in 100 mM increments. All buffers were pre-chilled on ice before use. Testicular cells (8.0×106) from adult WT mice were washed twice with ice-cold PBS and resuspended in Buffer A (0.3 M sucrose, 60 mM KCl, 60 mM Tris pH 8.0, 2 mM EDTA, 0.5% NP-40) supplemented with protease inhibitors. After 10 min rotation at 4 °C, nuclei were pelleted at 6000 ×g for 5 min at 4 °C. Each nuclear pellet was homogenized in 200 µL of 0 mM NaCl mRIPA with protease inhibitors by pipetting 15 times and incubated on ice for 3 min. Following centrifugation at 6500 ×g for 3 min at 4 °C, the supernatant was collected as the 0 mM fraction. This extraction was repeated sequentially with increasing NaCl concentrations. At salt concentrations exceeding 400 mM, the pellets became clear and highly viscous, failing to form a compact pellet; in such cases, the viscous material was collected from the tube lid after inversion. These samples were immediately used for subsequent analysis. Antibodies used in this study were summarized in Supplementary Table S3.

### Yeast two hybrid assay

The yeast two-hybrid assay was conducted by GeneCreate Biological Engineering (Wuhan, China). In brief, the full-length cDNA of mouse *Zcwpw2* was subcloned into the pGBKT7 vector to serve as the bait construct. Meanwhile, the full-length cDNAs of mouse *Iho1*, *Mei4*, *Rec114*, and *Hormad1* were separately inserted into the pGADT7 vector as prey constructs. The bait and prey plasmids were co-transformed into yeast two-hybrid Gold strain cells, and positive transformants were screened on selective nutrient-deficient medium (SD/-Leu/-Trp/-His).

### ChIP-seq library preparation and sequencing

The ChIP-seq libraries were prepared as previously described [49] with further modifications primarily for DNA purification. In brief, 2×10^6^ cells from PD13–PD14 testes were cross-linked in 100 μL of 1% formaldehyde in PBS at room temperature for 10 min, and then quenched with 25 μL 1.25M glycine solution and washed with PBS. The cells were then incubated in 150 μL lysis buffer (50 mM Tris-HCl pH 8.0, 10 mM EDTA pH8.0, 0.5% SDS, 1mM PMSF, and 1× proteinase inhibitor cocktail) for 20 min on ice then sonicated using a Diagenode Bioruptor sonication device for 23 cycles (30s ON and 30s OFF). A total of 150µl 300mM SDS-free RIPA buffer (10 mM Tris-HCl pH 7.5, 300 mM NaCl, 1 mM EDTA, 0.5 mM EGTA, 1% Triton X-100, 0.1% Na-deoxycholate, 1mM PMSF, 1×proteinase inhibitor cocktail, and 20 mM Na-butyrate) and 200µl 140mM SDS-free RIPA buffer was added to the samples. After centrifugation at 13,000 × g for 10 min at 4°C, 40 μL of the supernatant was removed and used as the sample input. The remaining supernatant was transferred to a 1 ml tube containing suspended antibody-coated Protein A beads, followed by incubation on a tube rotator overnight at 4°C. Next, the incubated Protein A beads were washed once with RIPA buffer containing 250 mM NaCl, washed three times with RIPA buffer containing 500 mM NaCl, and washed once with TE buffer (10 mM Tris-HCl pH 8.0, 1mM EDTA). Next, the beads were transferred to a new 0.5ml tube, followed by incubation in 100 μL ChIP elution buffer (10mM Tris-HCl pH8.0, 5mM EDTA, 300mM NaCl, and 0.5% SDS) containing 5 µL proteinase K (Qiagen, 20mg/ml stock) at 55°C for 2 h, and then at 65°C for 4 h. The eluate was transferred to a fresh 0.5 mL tube, and the enriched DNA was purified by 1.8X SPRIselect beads, followed by dissolution in 50 μL TE buffer. Finally, the NEBNext Ultra II DNA Library Prep Kit for Illumina (NEB, E7645S) was used for library construction according to the product instructions. Libraries were sequenced using the Illumina X-ten and NovaSeq 6000 platform in PE150 mode (Novogene, Beijing, China).

### End-seq library preparation and sequencing

END-seq was conducted following the protocol described by André et al[51]. In brief, mouse testes were dissociated into single cells and subsequently embedded in agarose. Immediately after solidification, the agarose-embedded cells were lysed and subjected to Proteinase K digestion. The resulting plugs were washed with TE buffer, treated with RNase, and stored at 4°C for up to one week prior to subsequent enzymatic processing. Following protein and RNA removal, DNA ends at DSBs were blunted using single-strand-specific exonucleases, then A-tailed and ligated within the agarose plugs to a biotin-labeled, T-tailed DNA adapter compatible with Illumina sequencing.

### ChIP-seq data analysis

ChIP-seq reads were processed by trimming all raw reads to 100 bp and removing low-quality reads with Trimmomatic (v0.32)[52]. Filtered paired-end reads were mapped to the mm10 mouse genome assembly using Bowtie2 (v2.3.4.2)[53] with the options “-X 2000 --no-discordant --no-contain”. Alignments with MAPQ values below 10 were discarded, and duplicate fragments introduced during PCR amplification were removed using Samtools[54] and Picard (https://broadinstitute.github.io/picard/). For peak detection, Reads from two replicate experiments were pooled and analyzed with MACS2 (v2.2.6)[55]. ZCWPW2 peaks were called with the parameters “--SPMR --nomodel -p 0.002”. Genome-wide enrichment tracks for ZCWPW2 was computed as ChIP-versus-input fold change with macs2 bdgcmp, followed by conversion of bedGraph output files into BigWig format using bedGraphToBigWig. The final ChIP-seq signal tracks were visualized by Integrative Genomics Viewer (IGV)[56]. To assess signal distribution around peaks or other regions, DeepTools[57] computeMatrix was applied to calculate normalized read density in 40-bp bins within a ±2-kb or ± 3-kb window centered on each peak center. The resulting signal matrices were visualized as heatmaps and average profiles using DeepTools (plotHeatmap, plotProfile) and R (v4.2.3). The findMotifsGenome.pl script in HOMER (v5.1)[58] was used to perform transcription factor motif enrichment analysis. The mm10 refGene and genomic annotation files, including promoter regions defined as TSS ± 2 kb, were downloaded from the UCSC Table Browser. BEDTools (v2.27.1)[59] was used to identify overlaps between peaks or genomic regions. Other processed data, including peak files and BigWig files for DMC1, PRDM9, H3K4me3, and H3K36me3, were either directly obtained from previously published studies and public datasets or processed using the same analytical pipeline, as summarized in Supplementary Table S2.

### End-seq data analysis

The high-quality reads aligned to the mouse genome were processed in a manner similar to the steps used in ChIP-seq data analysis. End-seq peaks were called using MACS2 with the parameters: “--SPMR --nomodel --extsize 2000 --shift -1000 --nolambda”. Genome-wide normalized End-seq signal tracks, representing DSBs, were generated using the bamCoverage tool from DeepTools with the parameters: “--normalizeUsing RPKM --binSize 20”. The resulting signal matrices were visualized as heatmaps and average profiles using DeepTools (plotHeatmap, plotProfile) and R (v4.2.3). Both the End-seq data generated in this study and the previously published End-seq data, as summarized in Supplementary Table S2, were analyzed.

### Statistics

Statistical analysis was carried out with GraphPad Prism 6. Unpaired *t*-tests were used to analyze differences between two groups. All tests and p-values are provided in the corresponding legends and/or figures. Quantification of fluorescence image data are summarized in Supplementary Table S4.

